# Coronary arterial development is regulated by a Dll4-Jag1-EphrinB2 signaling cascade

**DOI:** 10.1101/711333

**Authors:** Stanislao Igor Travisano, Vera Lucia Oliveira, Belén Prados, Joaquim Grego-Bessa, Vanesa Bou, Manuel José Gómez, Fátima Sánchez-Cabo, Donal MacGrogan, José Luis de la Pompa

## Abstract

Coronaries are essential for myocardial growth and heart function. Notch is crucial for mouse embryonic angiogenesis, but its role in coronary development remains uncertain. We show Jag1, Dll4 and activated Notch1 receptor expression in sinus venosus (SV) endocardium. Endocardial *Jag1* removal blocks SV capillary sprouting, while *Dll4* inactivation stimulates excessive capillary growth, suggesting that ligand antagonism regulates coronary primary plexus formation. Later endothelial ligand removal, or forced expression of Dll4 or the glycosyltransferase MFng, blocks coronary plexus remodeling, arterial differentiation, and perivascular cell maturation. Endocardial deletion of *Efnb2* phenocopies the coronary arterial defects of Notch mutants. Angiogenic rescue experiments in ventricular explants, or in primary human endothelial cells, indicate that EphrinB2 is a critical effector of antagonistic Dll4 and Jag1 functions in arterial morphogenesis. Thus, coronary arterial precursors are specified in the SV prior to primary coronary plexus formation and subsequent arterial differentiation depends on a Dll4-Jag1-EphrinB2 signaling cascade.

## Introduction

Coronary artery disease leading to cardiac muscle ischemia is the major cause of morbidity and death worldwide (Sanchis-Gomar et al. 2016). Deciphering the molecular pathways driving progenitor cell deployment during coronary angiogenesis could inspire cell-based solutions for revascularization following ischemic heart disease. The coronary endothelium in mouse derives from at least two complementary progenitor sources (Tian et al. 2015) that may share a common developmental origin (Zhang et al. 2016a; Zhang et al. 2016b). The sinus venosus (SV) commits progenitors to arteries and veins of the outer myocardial wall (Red-Horse et al. 2010; Tian et al. 2013), and the endocardium contributes to arteries of the inner myocardial wall and septum (Red-Horse et al. 2010; Wu et al. 2012; Tian et al. 2013). Regardless of origin, the endothelial precursors invest the myocardial wall along stereotyped routes and eventually interlink in a highly coordinated fashion. Subsequently, discrete components of the primitive plexus are remodeled into arteries and veins, and stabilized through mural cell investment and smooth muscle cell differentiation (Udan et al. 2013). Arterial-venous specification of endothelial progenitors is genetically pre-determined (Swift and Weinstein 2009), whereas arterial differentiation and patterning depend on environmental cues, such as blood flow and hypoxia-dependent proangiogenic signals (le Noble et al. 2005; Jones et al. 2006; Fish and Wythe 2015). Vascular endothelial growth factor (VEGF) binds endothelial receptors and drives the expansion of the blood vessel network as a response to hypoxia (Dor et al. 2001; Liao and Johnson 2007; Krock et al. 2011).

The Notch signaling pathway is involved in angiogenesis in the mouse embryo and in the postnatal retina. In this processes, the ligand Dll4 is upregulated by VEGF, leading to Notch activation in adjacent endothelial cells (ECs), vessel growth attenuation, and maintenance of vascular integrity (Blanco and Gerhardt 2013). In contrast, the ligand Jag1 has a proangiogenic Dll4-Notch–inhibitory function, suggesting that the overall response of ECs to VEGF is mediated by the opposing roles of Dll4 and Jag1 (Benedito et al. 2009). Dll4-Notch1 signaling is strengthened in the presence of the glycosyltransferase MFng (Benedito et al. 2009; D’Amato et al. 2016).

Our understanding of how coronary vessels originate, are patterned, and integrate with the systemic circulation to become functional is still limited. Coronary arteries are distinct from peripheral arteries in that they originate from SV and ventricular endocardium, which is a specialized endothelium that lines the myocardium. Moreover, SV endothelium has a venous identity, as opposed to the retina vascular bed, which has no pre-determined venous identity. Given these differences of developmental context, it is essential to evaluate the role of Notch in coronary arterial development, and importantly, the implications for heart development and repair.

Several components of the Notch pathway have been examined in the context of coronary artery formation. Inactivation of the Notch modifier *Pofut1* results in excessive coronary angiogenic cell proliferation and plexus formation (Wang et al. 2017), while endothelial inactivation of *Adam10*, required for Notch signaling activation, leads to defective coronary arterial differentiation (Farber et al. 2019). Transcriptomics has shown that pre-artery cells appear in the immature coronary vessel plexus before coronary blood flow onset, and express Notch genes, including *Dll4* (Su et al. 2018). Here, we examine the early and late requirements of Notch ligands Jag1 and Dll4, and their downstream effector EphrinB2, for coronary arterial development.

## Methods

### Mouse strains and genotyping

Animal studies were approved by the CNIC Animal Experimentation Ethics Committee and by the Madrid regional government (Ref. PROEX 118/15). All animal procedures conformed to EU Directive 2010/63EU and Recommendation 2007/526/EC regarding the protection of animals used for experimental and other scientific purposes, enforced in Spanish law under Real Decreto 1201/2005. Mouse strains were CBF1:H2B-Venus (Nowotschin et al. 2013), *Tie2-Cre* (Kisanuki et al. 2001), N*fatc1-Cre* (Wu et al. 2012), *Pdgfb-iCre^ERT2^*(Wang et al. 2010), *Cdh5-Cre^ERT2^, Dll4^flox^* (Koch et al. 2008), *Jag1^flox^*(Mancini et al. 2005), *Efnb2^flox^*(Grunwald et al. 2004), *Notch1^flox^* (Radtke et al. 1999), *Mfng^GOF^* (). To generate the *R26CAGDll4^GOF^* transgenic line, a full-length mouse *Dll4* cDNA was obtained from clone IMAGE 6825525. The sequence was PCR amplified with Phusion High-Fidelity DNA Polymerase (NEB) and primers containing *BamH*I and *Cla*I sites and was cloned in-frame with 6 Myc Tag epitopes into the *BamH*I and *Cla*I sites of a *pCS2-MT* plamid. The *Dll4*-MT fragment was subcloned into pCDNA3.1 with *BamH*I and *EcoR*I and excised with *BamH*I and *Not*I. We then modified *pCCALL2* (Lobe et al. 1999) by cloning new *Xba*I sites before and after a CAG cassette, which includes a CMV enhancer/*β-actin* promoter and a rabbit *β*-globin polyA signal. The *Kpn*I and *Not*I PCR *Dll4*-MT was cloned into the *Bgl*II-*Not*I sites of the modified *pCCALL2*. The *Xba*I-cassette containing *CAG-loxP-β-Geo-loxP-Dll4-IRESeGFP* was obtained by digestion and cloned into the *Xba*I site of the *pROSA26-1* plasmid (Soriano 1999). The final construct was linearized with *Xho*I and electroporated into R1 mESCs derived from a cross of 129/Sv x 129/Sv-CP mice (Nagy et al. 1993). After G418 (200 μg/ml) selection for 7 days, 231 clones were picked. Homologous recombination was identified by Southern blot of *EcoR*V-digested DNA and hybridized with 5’ and 3’ probes. About 25% of the clones were positive, and we selected three clones to confirm karyotype. One positive clone was injected into C57/BL6C blastocysts to generate chimaeras that transmitted the transgene to their offspring. The resulting founders were genotyped by PCR of tail genomic DNA using primers targeting the R26 locus before and after the cloning site and the transgene polyA signal (see Fig.4).

Throughout the MS, *Jag1^flox/flox^;Nfatc1-Cre* mice are called *Jag1^flox^;Nfatc1-Cre* for simplicity; this also applies to the *Dll4^flox^;Nfatc1-Cre* mice, *Jag1^flox^;Pdgfb-iCre^ERT2^* mice, *Dll4^flox^*;*Pdgfb-iCre^ERT2^, Dll4^flox^;Cdh5-Cre^ERTT2^, Jag1^flox^;Cdh5-Cre^ERT2^* mice, *Dll4^flox^*;*Cdh5-Cre^ERT2^, Dll4^flox^;Cdh5-Cre^ERTT2^MFng^GOF^;Tie2-Cre* mice (which carry two extra copies of *MFng*), and *Efnb2^flox^;Nfatc1-Cre* mice, but not to *Dll4^GOF^;Tie2-Cre* mice, which carry only one extra copy of *Dll4*.

### 4-hydroxy-tamoxifen (4-OHT) induction

Cre-inducible female mice were crossed with floxed Jag1 or Dll4 males, and pregnant females were administered with 4-OHT (H6278 Sigma) once by oral gavage at indicated gestation days (E9.5, 12.5 or E14.5). Embryos were dissected at E15.5 or E16.5 as indicated.

### Histology and *in situ* hybridization

Hematoxylin & eosin (H&E) staining and *in situ* hybridization (ISH) on sections were performed as described (Kanzler et al. 1998). Details of probes will be provided on request.

### Whole-mount immunofluorescence

Embryos were dissected and fixed for 2h in 4% paraformaldehyde (PFA) in PBS at 4°C, then permeabilized for 1h with 0.5% Triton X-100/PBS and subsequently blocked for 1h in Histoblock solution (5% goat serum, 3% BSA, 0.3% Tween-20 in PBS). After several washes in PBS-T (PBS containing 0.1% Tween-20), embryos were mounted in 1% agar in a 60mm petri dish. For whole-mount immunofluorescence on E11.5-E12.5 *CBF:H2B-Venus* embryos endogenous GFP, was imaged with a NIKON A1R confocal microscope. Z-stacks were captured every 5µm. For EdU immunofluorescence, pregnant females mice at E12.5 were injected intraperitoneally with 100 µl EdU nucleotides (2 mg ml−1 in PBS). The mice were euthanized 1 h later, and EdU incorporation was detected with the Click-iT EdU Imaging Kit (Thermo Fisher Scientific, C10340). For IsoB4 and Endomucin immunostaining on E11-4-E16.5 embryos, imaging by confocal z-projection of the deeper section of the myocardium captures the arteries (IsoB4-stained) with the outer section corresponding to the veins (IsoB4+Endomucin-stained). N1ICD embryos were fixed for 5h at 4°C in frozen Methanol, washed and incubated 20min at 98oC in DAKO target retrieval solution pH6 (Agilent). After washing in H_2_O, hearts were fixed again in acetone at -20°C for 10 min. After washing in PBT the hearts were blocked in 10% FBS, 5%BSA, 0.4% TritonX100 for 3h at RT under gentle rocking. Hearts were incubated with primary N1ICD-antibody at 1:500 for 2 days at 4°C and allowed to settle at RT or 1h. This was followed by washing 5h in PBS/0.4% TrtonX100 at RT. Hearts were then incubated in presence of anti-rabbit HRP (1:500), DAPI (1:1500) and anti VEcadherin (1:500) overnight at 4°C. After resting for 1h at RT, hearts were washed in PBT, and 30 min in presence TSA (tyramides) diluted 1:200 in PBS. Quantifications were carried out using ImageJ software. The main coronary trees were selected with the tool “Freehand selection” manually and measured directly from the autoscaled images obtained by Z-projection. The selected area was quantified in µm^2. Confocal microscopy analysis was carried out on a Nikon A1R or Leica LAS-AF 2.7.3.

### Immunofluorescence on sections

Paraffin sections (7µm) were incubated overnight with primary antibodies, followed by 1h incubation with a fluorescent-dye-conjugated secondary antibody. N1ICD, Dll4, Jag1, and p27 were immunostained using tyramide signal amplification (TSA) (Del Monte et al. 2007); see Supplemental Table 4 for antibodies. For p27, N1ICD SMA and Notch3 quantification, the number of positive nuclei was divided by the total number of nuclei counted on sections (≥3). The number of mural cells surrounding intramyocardial vessels expressing both SMA+ and Notch3+ (double positives) was counted and divided by the total number of intramyocardial vessels defined by Iso B4 staining in the same section. The number of subepicardial SMA+, Notch3+ double-positive cells was counted and divided by the total number of subepicardial vessels defined by IsoB4. Images were processed with ImageJ software.

### Mouse ventricular explant culture and immunofluorescence

Heart ventricular explants were performed as previously described with minor modifications (Wu et al. 2012). Ventricles were dissected from E10.5 or E11.5 embryos (with removal of the outflow tract and atria), rinsed with PBS to remove circulating cells, and placed in Nunc 4-well plates. Matrigel (Corning® Matrigel® Basement Membrane Matrix, *LDEV-Free, 10mL growth factor reduced, BD Biosciences 354234) was diluted 1:1 with DMEM plus 10% FBS and 10 ng/ml Vegf (Human-Vegf 165 Peprotech). Each well had a total volume of 400µl, and 3-4 hearts were cultured for 6 days and then fixed in 4% PFA4% 10 minutes and washed twice with PBS1X for 15minutes. Explants were permeabilized with 0.5% Triton X-100 for 1 hour, blocked with Histoblock solution (FBS Histoblock) for 2-3 hours at RT, and incubated with anti-CD31 1:100 (Purified Rat Anti-Mouse CD31 BD Biosciences Pharmingen) overnight at 4°C. Anti-rat biotinylated was used as secondary antibody and diluted in BSA 1:150. Staining was amplified with the ABC kit and 3 minutez of TSA (tyramides) diluted 1:100 in PBS.

### Quantification of compact myocardium thickness

The method used was a modification of that described by (Chen et al. 2009; Yang et al. 2012). Briefly, 7µm paraffin sections from E12.5, E15.5 and E16.5 wild type (WT) and mutant hearts were stained with anti-CD31 or Endomucin (emcn) and anti-MF20 antibodies to visualize ventricular structures. Confocal images were obtained with a NIKON A1R confocal microscope. Measurements were made using ImageJ software. In E15.5 and E16.5 heart sections, endocardial cells were stained with anti-emcn and myocardium with anti-cTnT. Left and right ventricles were analyzed separately. For each measurement, settings were kept constant for all images, using the scale bar recorded in each image as the reference distance. The thickness of the compact myocardium was measured by dividing the ventricle into left and right parts. Several measurements were taken in each region and the mean was expressed in µm.

### India ink perfusion

Embryos were collected from E15.5 to E18.5 in PBS containing 2µg/ml heparin. The thoracic cavities were immediately cut and transferred to DMEM containing 10%FBS and heparin on ice. Hearts were carefully dissected from the surrounding tissue and then placed on an inverted Petri dish and gently dried to avoid movement during perfusion. India ink or red tempera was diluted in PSB/heparin and injected from the ascending aorta with a borosilicate glass tube thinned to the appropriate diameter and attached to a mouth pipette. India ink was slowly injected by minute puffs of breath during diastolic intervals. Hearts were fixed in 4% PFA, dehydrated, cleared in BABB (benzyl alcohol: benzyl benzoate, 1:1) and imaged with a stereomicroscope.

### Lentiviral production

Bacterial glycerol stocks for JAG1 and DLL4 MISSION shRNA were purchased from SIGMA. MISSION shRNAs: shRNA JAGGED1_2: clone ID: NM_000214.2-3357s21c1; shRNA JAGGED1_3: clone ID: NM_000214.2-1686s21c1; shRNA DLL4_1: clone ID_ NM_019074.2-2149s21c1; shRNA DLL4_1: clone ID_ NM_019074.2-821s21c1. Concentrated lentiviral particles were produced by triple transfection in HEK293T cells. Briefly, HEK293T cells were cultured in 150-mm plates for transfection of the lentiviral vectors. The psPAX2 packing plasmid, the pMD.G envelope plasmid coding for the VSV-G glycoprotein (Viral Vectors Unit, CNIC, Spain) and the LTR-bearing shuttle lentiviral plasmids were cotransfected using the calcium phosphate method. The transfection medium was replaced with fresh medium 16 hours post-transfection. Viral supernatants were harvested at 72 hours post-transfection, filtered through a 0.45-µm filter (Steriflip-HV, Millipore, MA, USA) and concentrated by ultracentrifugation. Precipitated viruses were resuspended in pre-chilled 1X PBS, aliquoted and stored at -80° C for further use. Viral titer was measured on viral genomes by qPCR using the standard curve method with primers against the LTRs. (Forward primer:5’-AGCTTGCCTTGAGTGCTTCAA-3’;Reverse primer:5’-AGGGTCTGAGGGATCTCTAGTTA-3’).

Full-length murine *EphrinB2 (Efnb2)*, kindly provided by Ralf Adams (MPI for Molecular Biomedicine, University of Münster, Germany), was subcloned into the lentiviral vector pRLL-IRES-eGFP (Addgene). Concentrated lentiviruses expressing pRLL-IRES-eGFP or pRLL-MFng-IRES-eGFP were obtained as described (Esteban et al. 2011). Viruses were titrated in Jurkat cells, and infection efficiency (GFP-expressing cells) and cell death (propidium iodide staining) were monitored by flow cytometry.

### Culture and infection on HUVEC

Human umbilical vein endothelial cells (HUVECs) were maintained in supplemented EGM2 medium (EGM2 Bulletkit, K3CC-3162 Lonza). Cells were transduced on suspension at a multiplicity of infection (m.o.i.) of 80 with combinations of 2 shRNAs against JAG1 or DLL4 together with GFP or EFNB2 overexpressing lentivirus and seeded onto 24-well plates or 12-well plates at a density of 3×104 or 6×104 cells/well respectively. Infection efficiency was near 100%. HUVECs were harvested after 48h. Gene expression analysis was performed using KiCqStart SYBR Green predesigned primers (Sigma-Aldrich).

### Angiogenesis assays on Matrigel

The remodeling stage of angiogenesis was investigated using endothelial cell tube formation assay on Matrigel (#354234 Corning). Briefly 50μl of ice cold Matrigel was coated on a 96 micro-well plate as a base for tube formation. After allowing the gel to settle in the incubator for 30 min at 37 °C, 5% CO2, HUVECs in basic EGM medium (EBM2 H3CC-3156, Lonza) were seeded in triplicate at a density of 12.5 x104 on Matrigel and incubated at 37 °C, 5% CO2. After 5h tube formation was monitored by phase-contrast microscope. The tube networks were quantified using NIH Image J with Angiogenesis plugin software (http://image.bio.methods.free.fr/ImageJ/?Angiogenesis-Analyzer-for-ImageJ). Excel data were represented on Graphpad Prism 7 as mean with SD. For statistical analysis a one-way ANOVA and Tukey post-hoc tests were performed. P <0.05 was considered significant.

Angiogenesis *in vitro* was investigated using endothelial cell tube formation assay on Matrigel as previously described (Grant et al. 1991). The tube networks were quantified using NIH Image J with Angiogenesis plugin software http://image.bio.methods.free.fr/ImageJ/?Angiogenesis-Analyzer-for-ImageJ.

### Statistical analysis

Statistical analysis was carried out using Prism 7 (GraphPad). All statistical tests were performed using two-sided, unpaired Student’s t-tests, except for Figure 1 and Figure 2 where we performed one-way ANOVA with Tukey’s multiple comparison tests to assess statistical significance with a 95% confidence interval, where numerical data are presented as mean ±SD; results are marked with one asterisk (*) if P <0.05, two (**) if P <0.01 and three (***) if P <0.001. Sample size was chosen empirically according to previous experience in the calculation of experimental variability. No statistical method was used to predetermine sample size. All experiments were carried out with at least three biological replicates. The numbers of animals used are described in the corresponding figure legends. Animals were genotyped before the experiment and were caged together and treated in the same way. Variance was comparable between groups throughout the manuscript. We chose the appropriate tests according to the data distributions. The experiments were not randomized. The investigators were not blinded to allocation during experiments and outcome assessment.

**Figure 1.**
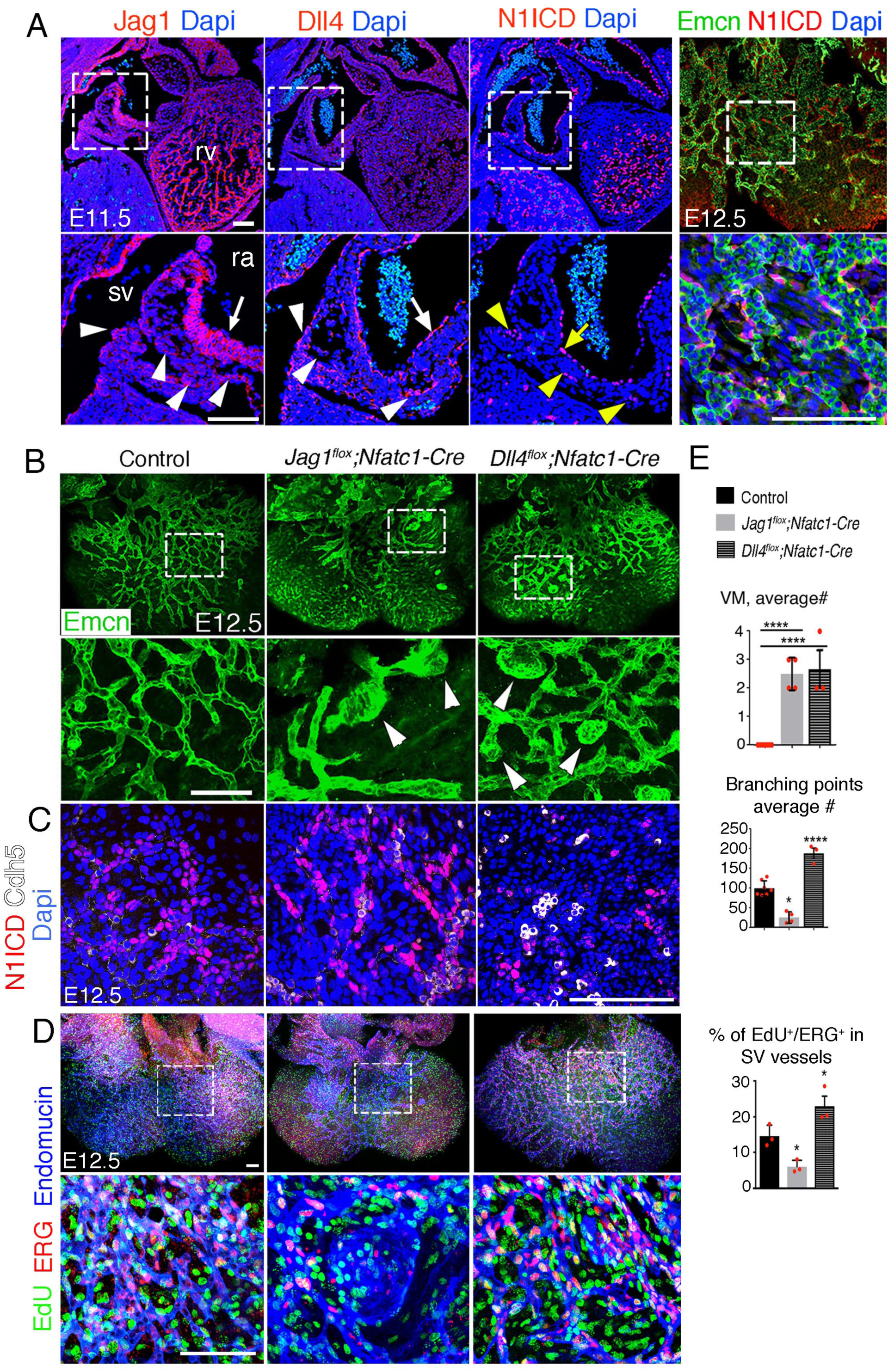
Endocardial *Jag1* or *Dll4* inactivation disrupts coronary plexus formation. (**A**), Jag1, Dll4, and N1ICD immunostaining (red) in E11.5 control hearts, sagittal views. Magnified views of boxed areas show details of sinus venosus (sv, arrowheads) and right atrium (ra, arrow). Whole-mount dorsal view of immunostainings for N1ICD (red) and Emcn (green) in E12.5 control heart. Magnified views show detail of sub-epicardial endothelium. Nuclei are counterstained with Dapi (blue). (**B**) Whole-mount dorsal view of immunostaining for Emcn (green) in E12.5 control, *Jag1^flox^;Nfatc1-Cre*), and *Dll4^flox^;Nfatc1-Cre* mutant hearts. Arrowheads indicate vascular malformations. Quantified data of average number of vascular malformations (VM) and average number of branching points in E12.5 control, *Jag1^flox^;Nfatc1-Cre* and *Dll4^flox^;Nfatc1-Cre* hearts. (**C**) Dorsal views of whole-mount E12.5 control, *Jag1^flox^;Nfat-Cre*, and *Dll4^flox^;Nfatc1-Cre* hearts stained for N1ICD (red) and VE-Caderin (white). Microscope: Leica SP5. Software: LAS-AF 2.7.3. build 9723. Objective: HCX PL APO CS 10x 0.4 dry. HCX PL APO lambda blue 20x 0.7 multi-immersion. (**D**) Dorsal views of whole-mount E12.5 control, *Jag1^flox^;Nfat-Cre*, and *Dll4^flox^;Nfatc1-Cre* hearts stained for EdU (green), ERG (red), and Emcn (blue). Scale bars, 100 µm. Microscope: Nikon A1-R. Software: NIS Elements AR 4.30.02. Build 1053 LO, 64 bits. Objectives: Plan Apo VC 20x/0.75 DIC N2 dry; Plan Fluor 40x/1.3 Oil DIC H N2 Oil. (**E**) Quantified data for vascular malformations (VM), average # of branching points and EdU-ERG dual-positive nuclei as a percentage of all nuclei in sub-epicardial vessels. Data are mean ± s.d. (n= 3-4 control embryos and n= 3-4 mutant embryos). \**P* < 0.05, \*\*\*\**P* < 0.0001 by Student’s *t*-test. Abbreviations: rv, right ventricle.

**Figure 2.**
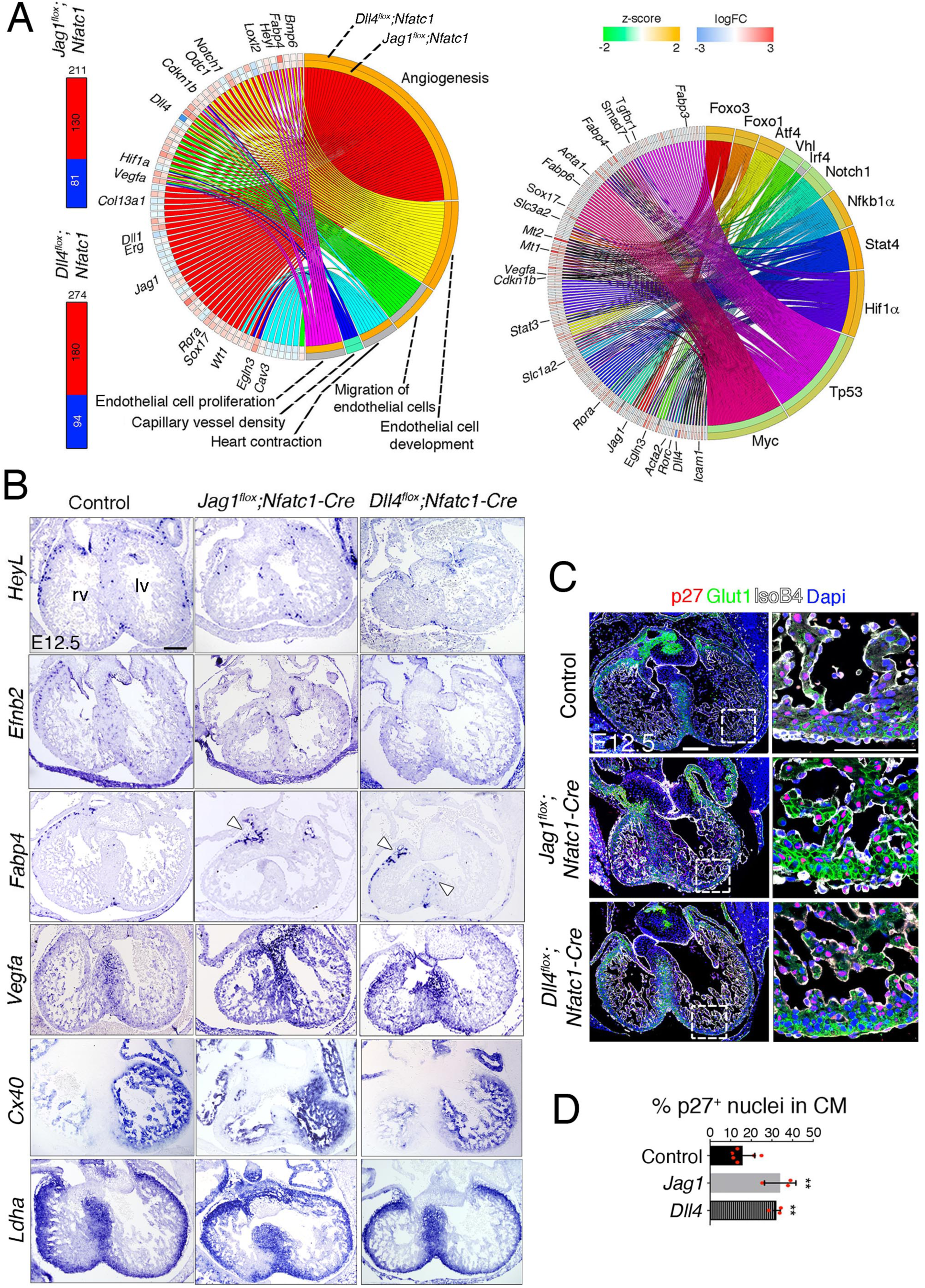
Transcriptome profiling of endocardial *Jag1* and *Dll4* mutant hearts. (**A**) *Left*, Total number of differentially expressed genes identified by RNA-seq (Benjamini-Hochberg (B-H) adjusted P < 0.05) in the indicated genotypes. Numbers indicate upregulated genes (red) and downregulated genes (blue). *Center*, circular plot of representative differentially expressed genes, presenting a detailed view of the relationships between expression changes (left semicircle perimeter) and IPA functions belonging to the Cardiovascular System Development and Function category (right semicircle perimeter). For both circular plots, in the left semicircle perimeter, the inner ring represents *Jag1^flox^;Nfatc1-Cre* data and the outer ring *Dll4^flox^;Nfatc1-Cre* data. Activation z-score scale: green, repression; orange, activation; white, unchanged. LogFC scale: red, upregulated; blue, downregulated; white, unchanged. Right, circular plot showing representative differentially expressed genes depending of selected upstream regulators. Details in Table supplement 2 and Table supplement 3. (**B**) *In situ* hybridization (ISH) of *HeyL, Efnb2*, *Fabp4, Vegfa, Cx40* and *Ldha* on E12.5 control, *Jag1^flox^;Nfatc1-Cre*, and *Dll4^flox^;Nfatc1-Cre* heart sections. Arrowheads indicate *Fabp4* expression in capillary vessels. (**C**) Immunohistochemistry of p27 (red), Glut1 (green), and IsoB4 (white) on E12.5 control, *Jag1^flox^;Nfatc1-Cre*, and *Dll4^flox^;Nfatc1-Cre* mutant heart sections. Dapi counterstain (blue). Microscope: Nikon A1-R. Software: NIS Elements AR 4.30.02. Build 1053 LO, 64 bits. Objectives: Plan Apo VC 20x/0,75 DIC N2 dry; Plan Fluor 40x/1,3 Oil DIC H N2 Oil. Quantified data for p27-positive nuclei as a % of total CM nuclei. Data are mean ± s.d. (n= 3 sections from 4-7 control embryos and n= 3 from 3-5 mutant embryos). \*\**P* < 0.01 by Student’s *t*-test. Abbreviations: lv, left ventricle; rv, right ventricle. Scale bars, 100 µm.

### RNA-sequencing

RNA was isolated at E12.5, from whole hearts of *Jag1^flox^;Nfatc1-Cre* and *Dll4^flox^;Nfatc1-Cre* embryos, as well as their WT counterparts (4 replicate samples for each condition, pooling 4 embryos per sample, in all cases). At E15.5, RNA was isolated from ventricles of *Dll4^flox^;Cdh5-Cre^ERT2^* (3 replicate samples, pooling 3 embryos per sample), as well as their WT counterparts (4 replicate samples, pooling 3 embryos per sample). RNA-Seq data for *Jag1^flox^;Nfatc1-Cre*, *Dll4^flox^;Nfatc1-Cre*, and *Dll4^flox^;Cdh5-Cre^ERT2^*experiments was generated by CNIC’s Genomics Unit. RNA-Seq sequencing reads were pre-processed by means of a pipeline that used FastQC (http://www.bioinformatics.babraham.ac.uk/projects/fastqc/) to assess read quality and Cutadapt v1.6 (Martin 2011) to trim sequencing reads, thus eliminating Illumina adaptor remains, and to discard reads shorter than 30 bp. Resulting reads were mapped against the mouse transcriptome (GRCm38 assembly, Ensembl release 76) and quantified using RSEM v1.2.20 (Li and Dewey 2011). Around 80-90% of the reads participated in at least one reported alignment. Expected expression counts calculated with RSEM were then processed with an analysis pipeline that used Bioconductor package Limma (Ritchie et al. 2015) for normalization and differential expression testing in six pairwise contrasts involving mutant versus WT comparisons. Changes in gene expression were considered significant if associated with a Benjamini-Hochberg (BH) adjusted P-value < 0.05. The number of differentially expressed genes detected in comparisons between *Jag1^flox^;Nfatc1-Cre*, *Dll4^flox^;Nfatc1-Cre* and *Dll4^flox^;Cdh5-Cre^ERT2^* and their control counterparts was 205, 257 and 156, respectively. Enrichment analyzes were performed with IPA (Ingenuity Pathway Analysis, Quiagen, http://www.ingenuity.com). IPA was used to identify collections of genes associated with canonical pathways, common upstream regulators, or functional terms significantly overrepresented in the sets of differentially expressed genes; statistical significance was defined by Benjamini-Hochberg adjusted P-value <0.05. Circular plots summarizing the association between genes and enriched pathways, upstream regulators, and functional terms were generated with GOplot (Walter et al. 2015).

### Accession number

Data are deposited in the NCBI GEO database under accession number GSE110614. The following secure token has been created to allow review of record GSE110614 while it maintains private status: czgniocarzqzrcr

## Results

### Jag1 and Dll4 are expressed in SV endocardium and coronary vessels endothelium

We examined SV and coronary vessels for the expression of Jag1 and Dll4. At embryonic day 11.5 (E11.5), Jag1 was detected in SV ECs and in ECs extending into the right atrium (Figure 1A). Dll4 was also expressed in ECs emanating from the SV and in the endocardium lining the right atrium (Figure 1A). These results suggest that either ligand could potentially activate Notch1 in the SV endocardium (Figure 1A). At E12.5, N1ICD was detected in endomucin (Emcn)-positive ECs in subepicardial capillaries emerging from the SV (Figure 1A). At E13.5, Jag1 and *Dll4*, were expressed in ECs of the developing coronary arteries (intramyocardial vessels; Supplemental Figure 1A). *Dll4* and *MFng* were also expressed in prospective veins (subepicardial vessels; Supplemental Figure 1A). At E15.5, Jag1, *Dll4*, and *MFng* were all expressed in arterial ECs, whereas *MFng* was still found in subepicardial vessels (Supplemental Figure 1B). Thus, Jag1 and Dll4 expression is found in discrete ECs in SV endothelium, abates in sub-epicardial veins and becomes restricted to intramyocardial coronary arteries at later developmental stages.

### Nfatc1^+^ progenitors give rise to the majority of subepicardial vessels

To inactivate Notch ligands in SV progenitors we used the *Nfatc1-Cre* driver line (Wu et al. 2012). To confirm the SV and endocardial specificity of this line, we crossed it with the *Rosa26-LacZ* reporter line (Soriano 1999). X-gal-staining of heart sections of E11.5 embryos identified patchy LacZ expression in SV endothelium (Supplemental Figure 2A). *β*-gal staining was consistent with Nfatc1 protein nuclear localization in a subset of ECs lining the SV (Supplemental Figure 2A). Uniform *β*-gal staining was detected in ventricular endocardium and cushion mesenchyme derived from endocardial cells (Supplemental Figure 2A). At E12.5, co-labelling with an anti-Pecam1 antibody revealed *β*-gal^+^ staining in 60% of Pecam1^+^ subepicardial vessels in the right ventricle and about 50% in the left ventricle (Supplemental Figure 2B). Tracking the expression of *Nfatc1-Cre*-driven red fluorescent protein (RFP) and the endothelial-specific nuclear protein Erg at E12.5 indicated that about 80% of nuclei in the endothelial network were Nfatc1^+^Erg^+^ (Supplemental Figure 2C). Thus, SV-derived Nfatc1^+^ progenitors give rise to 50-80% of subepicardial vessels in the ventricular wall, consistent with previous reports (Chen et al. 2014; Cavallero et al. 2015; Zhang et al. 2016a).

To trace the fate of Nfatc1^+^ cells relative to Notch activity, we crossed *Nfatc1-Cre;Rosa26-RFP* mice with the Notch reporter line *CBF:H2B-Venus*. At E11.5, a subset of nuclear-stained RFP ECs extending from the SV into the right ventricle were co-labelled with *CBF:H2B-Venus* (Supplemental Figure 2D). At E12.5, some RFP-labelled capillaries on the dorsal side of the heart were co-labelled with *CBF:H2B-Venus* while others were labelled with *CBF:H2B-Venus* alone (Supplemental Figure 2E), indicating that Notch signaling activity is present in both Nfatc1^+^ and Nfatc1^-^ populations of SV-derived ECs.

### Opposing roles of *Jag1* and *Dll4* in coronary plexus formation from SV

*Jag1* inactivation with the *Nfatc1-Cre* driver line, specific of SV and endocardium (see Supplemental Figure 2 and related text), resulted in death at E13.5 (Supplemental Table 1). Whole-mount Endomucin (Emcn) staining at E12.5 revealed a well-formed vascular network covering the dorsal aspect of control hearts (Figure 1B,E; Supplemental Figure 3A), whereas the vascular network in *Jag1^flox^;Nfatc1-Cre* mutants was poorly developed, and exhibited numerous capillary malformations (Figure 1B,E; Supplemental Figure 3B) with decreased endothelial branching (Figure 1B). N1ICD expression was also more prominent (Figure 1C) and EC proliferation reduced 60% relative to control (Figure 1D,E). Ventricular wall thickness in endocardial *Jag1* mutants was reduced by 50-60% relative to controls (Supplemental Figure 4A). Thus, endocardial *Jag1* deletion arrests primitive coronary plexus formation and myocardial growth.

Removal of *Dll4* with the *Nfatc1-Cre* driver resulted in the death of high proportion of embryos at E10.5 (Supplemental Table 1). Nonetheless, about a third of mutant embryos survived until E12.5 (Supplemental Table 1). Whole-mount Emcn immunostaining of E12.5 *Dll4^flox^;Nfatc1-Cre* hearts revealed a comparatively denser capillary network (Figure 1B,E), characterized by numerous capillary malformations (Figure1B and Supplemental Figure 3A,C), and increased endothelial branching (Figure 1B,E). N1ICD expression was reduced (Figure 1C), as expected, and EC proliferation increased 30% relative to control (Figure 1D,E). These defects were associated with 50-60% reduction in ventricular wall thickness (Supplemental Figure 4A). Thus, endocardial inactivation of *Dll4* causes excessive expansion of the primitive coronary plexus and impairs myocardial growth.

### Arrested myocardial growth in endocardial *Jag1* or *Dll4* mutants is due to a global hypoxic and metabolic stress response

To determine the effect of early endocardial *Jag1* or *Dll4* inactivation on cardiac development we performed RNA-seq. This analysis yielded 211 differentially expressed genes (DEG) in the *Jag1^flox^;Nfatc1-Cre* transcriptome (130 upregulated, 81 downregulated; Figure 2A; supplemental Table 2; supplemental sheet 1) and 274 DEGs in the *Dll4^flox^;Nfatc1-Cre* transcriptome (180 upregulated, 94 downregulated; Figure 2A; supplemental Table 2; supplemental sheet 2).

Ingenuity Pathway Analysis (IPA) identified enrichment of EC functions (Figure 2A left plot; supplemental Table 3). The main terms overrepresented in both genotypes were angiogenesis and EC development, and capillary vessel density, possibly reflecting the lack of a normal-sized capillary network (Figure 2A; supplemental Table 3). EC proliferation and heart contraction were predicted to be upregulated in *Jag1^flox^;Nfatc1-Cre* embryos, while EC migration was upregulated in *Dll4^flox^;Nfatc1-Cre* mice (Figure 2A; Supplemental Table 3, sheets 1 and 2). Analysis of upstream regulators revealed activation of hypoxia (Hif1*α*), acute inflammatory response (Nf-*κ*b1*α*), intracellular stress pathways (Atf4), and response to metabolic stress pathways (Foxo), while cell cycle and DNA repair pathways (Myc, Tp53) were negatively regulated (Figure 2A, right plot).

*In situ* hybridization (ISH) showed reduced expression of *HeyL* and the Notch target *Efnb2* in subepicardial vessels (Figure 2B). *Fabp4,* a member of the fatty acid binding protein family, was upregulated (Figure 2A). Fabp4 is a DLL4-NOTCH target downstream of VEGF and FOXO1 in human EC (Harjes et al. 2014), required for EC growth and branching (Elmasri et al. 2009). *Fapb4* expression was found exclusively in the atrio-ventricular groove in *Jag1^flox^;Nfatc1-Cre* hearts (Figure 2B), but extended sub-epicardially into the base of the heart in *Dll4^flox^;Nfatc1-cre* hearts (Figure 2B). *Vegfa* was upregulated in all cardiac tissues (Figure 2A,B), suggesting a cardiac hypoxic response in mutant embryos. Cell cycle-associated genes such as *Cdkn1b/p27*, a negative regulator of cell proliferation, were also upregulated in both genotypes (Figure 2A), and p27 nuclear staining was increased 2-fold throughout the heart (Figure 2C,D), indicating decreased cell proliferation. *Connexin 40* (*Cx40*) and *Hey2*, which label trabecular and compact myocardium respectively, showed no alteration in their expression domains by ISH (Figure 2B and Supplemental Figure 4B), suggesting that chamber patterning was normal in these mutants. Likewise, the glycolytic marker genes *Ldha* and *Pdk1* (Menendez-Montes et al. 2016) were confined, as normal, to the compact myocardium (Figure 2B and Supplemental Figure 4B), indicating maintenance of ventricular chamber metabolic identity.

Thus, endocardial deletion of *Jag1* or *Dll4* causes malformation of the primary coronary plexus. The resulting activation of hypoxia and cellular stress pathways in both genotypes lead to arrested myocardial growth, while ventricular patterning remains unaffected.

### Defective coronary remodeling and maturation in endothelial *Jag1* or *Dll4* mutants

We next examined the requirements of endothelial Jag1 and Dll4 for coronary vessel remodeling and maturation. To circumvent the early lethality of *Jag1^flox^*;*Nfatc1-Cre* mutants, we crossed *Jag1^flox^* mice with tamoxifen-inducible and vascular endothelium-specific *Pdgfb-iCre^ERT2^*transgenic mice (Wang et al. 2010) to obtain *Jag1^flox^*;*Pdgfb-iCre^ERT2^* mice. Tamoxifen-induced *Jag1* deletion at E12.5 resulted in complete absence of arteries at E15.5, whereas the veins appeared unaffected (Figure 3A,C). We measured NOTCH pathway activity by carrying out a N1ICD staining that showed a 50% increase in endothelial N1ICD (Figure 3B,C), consistent with Jag1 acting as an inhibitory Notch ligand. Next, we examined perivascular coverage of the *Jag1^flox^*;*Pdgfb-iCre^ERT2^*endothelial coronary tree. We used *α*-smooth muscle actin (*α*SMA) and Notch3 to measure the extent of coronary vessel smooth muscle cell differentiation and pericyte coverage, respectively (Volz et al. 2015). We found that the proportion of *α*SMA- and Notch3-positive cells was significantly reduced in E16.5 *Jag1^flox^*;*Pdgfb-iCre^ERT2^*coronary arteries (Figure 3B,C) reflecting the lack of differentiation of perivascular cells.

**Figure 3.**
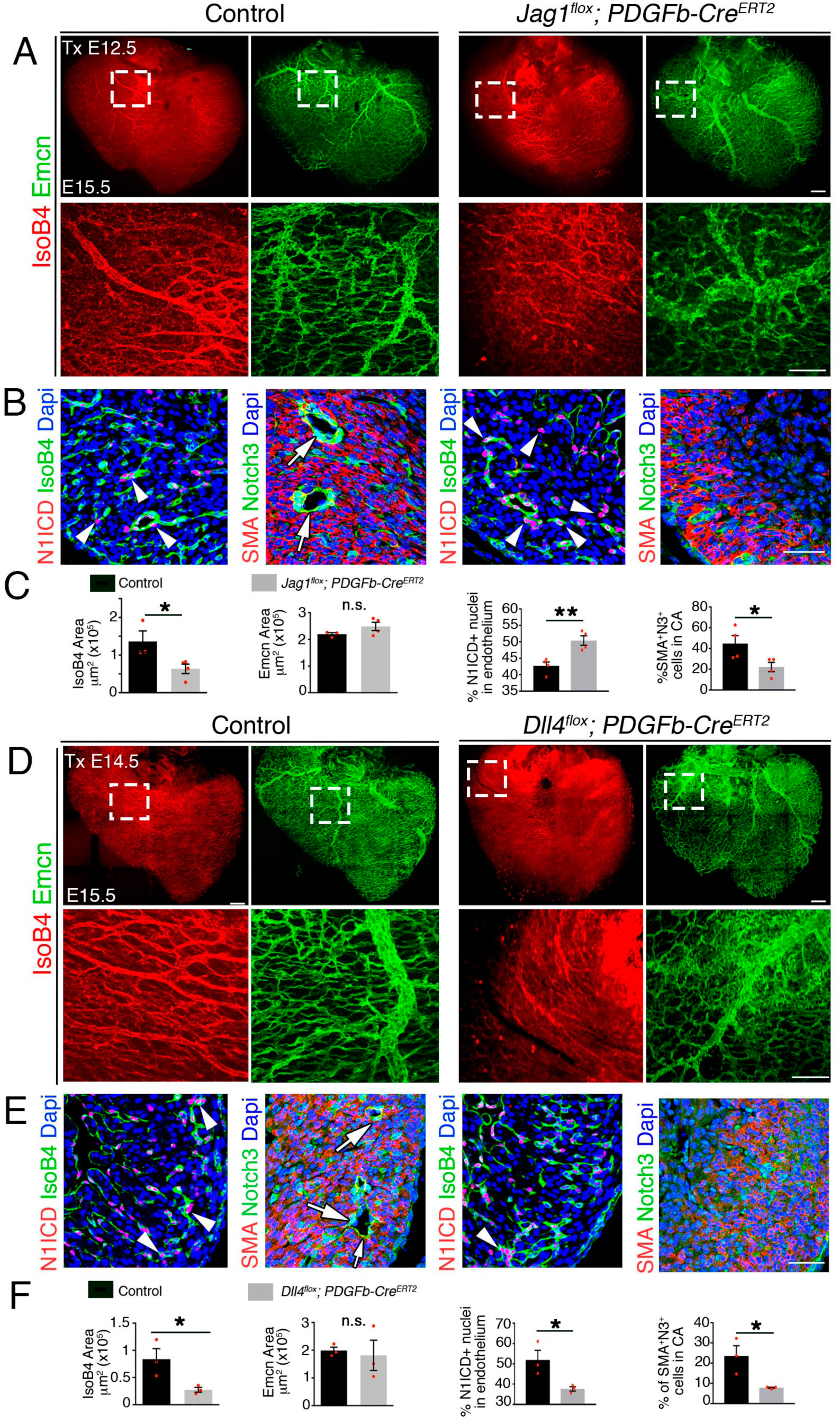
Late endothelial Jag1 or Dll4 inactivation disrupts coronary plexus remodeling. (**A**) Dorsal view of whole-mount immunochemistry for IsoB4 (red) and Emcn (green) in E15.5 control and *Jag1^flox^;Pdgfb-^iCreERT2^*mutant hearts, tamoxifen (Tx)-induced at E12.5. Scale bars, 100 µm. (**B**) E15.5 control and *Jag1^flox^; Pdgfb -iCre^ERT2^* mutant heart sections. Left, Immunostaining for N1ICD (red) and IsoB4 (green). Right, α-smooth-muscle actin (SMA, red) and Notch3 (green). Dapi counterstain (blue). Arrowheads point to N1ICD-positive nuclei. Arrows point to aSMA-Notch3 co-immunostaining. Yellow arrows point to small caliber intramyocardial vessels in the Jag1 mutant. Scale bars, 100 µm. (**C**) Quantified data from E15.5 control and *Jag1flox; Pdgfb -iCreERT2* hearts: area of coverage by coronary arteries and veins; N1ICD-positive nuclei as a percentage of total endothelial nuclei; and SMA-Notch3 co-immunostaining in coronary arteries. (**D**) Whole-mount dorsal view of immunohistochemistry for Emcn (green) and IsoB4 (red) in E15.5 control and *Dll4^flox^;Pdgfb-^iCre/ERT^* mutant hearts. Tx-induced at E14.5. (**E**) Left, immunohistochemistry for N1ICD (red) and IsoB4 (green); right, immunohistochemistry for SMA (red) and Notch3 (green) on control E15.5 WT and *Dll4^flox^; Pdgfb-iCre^ERT2^* mutant heart sections. Dapi counterstain (blue). Arrowheads indicate N1ICD-positive nuclei. Arrows point to SMA-Notch3 co-immunostaining. Yellow arrows point to small caliber intramyocardial vessels in the Dll4 mutant. Scale bars, 100 µm. Microscopes: Nikon A1-R, Leica SP5. Softwares: NIS Elements AR 4.30.02. Build 1053 LO, 64 bits (Nikon); LAS-AF 2.7.3. build 9723 (Leica). Objectives: Plan Apo VC 20x/0,75 DIC N2 dry; Plan Fluor 40x/1,3 Oil DIC H N2 Oil (Nikon); HCX PL APO CS 10x 0,4 dry. HCX PL APO lambda blue 20x 0,7 multi-immersion (Leica). (**F**) Quantified data from E16.5 control and *Dll4^flox^; Pdgfb -iCre^ERT2^* hearts: area covered by coronary arteries; area covered by veins; percentage of N1ICD-positive nuclei in endothelium as a percentage (%) of total nuclei; and SMA-Notch3 co-immunostaining in coronary arteries. Data are mean +/-s.d. (n= 3 sections from 2-4 Control embryos and from 2-4 mutant embryos). \**P* < 0.05, \*\**P* < 0.01, by Student’s t-test; n.s., not significant. Scale bars, 100 µm.

To examine Dll4 function in coronary artery formation we crossed *Dll4^fllox^* mice with *Pdgfb-iCre^ERT2^* to obtain *Dll4^flox^*;*Pdgfb-iCre^ERT2^*mice. Tamoxifen-induced *Dll4* deletion from E12.5 or E13.5, resulted in embryonic lethality at E15.5, confirming that embryo survival is highly sensitive to below normal endothelial Dll4 expression. However, induction at E14.5 resulted in the complete absence of arteries at E15.5, whereas veins were unaffected (Figure 3D,F). Endothelial N1ICD staining was decreased by 60% compared with controls (Figure 3E,F), consistent with Dll4 activating Notch1 during angiogenesis. Furthermore, *Dll4^flox^*;*Pdgfb-iCre^ERT2^*mutants had deficient perivascular cell coverage as indicated by decreased *α*SMA- and Notch3-positive cells (Figure 3E,F).

### Endocardium/endothelial Notch ligand inactivation impairs coronary artery formation and ventricular growth

To confirm endothelial Jag1 requirement for coronary arterial formation we crossed *Jag1^flox^*mice with *Cdh5-Cre^ERT2^* driver line (Wang et al. 2010) to obtain *Jag1^flox^;Cdh5-Cre^ERT2^* mice. Tamoxifen-induced *Jag1* inactivation from E9.5 onwards (Supplemental Figure 5A) resulted in reduced coronary artery coverage and marginally increased vein coverage (Supplemental Figure 5A,C). N1ICD staining in the endothelium was increased (Supplemental Figure 5B,C), reflecting Jag1 inhibitory Notch function during angiogenesis. Furthermore, *Jag1^flox^;Cdh5-Cre^ERT2^*mutant heart coronaries had decreased perivascular cell coverage as indicated by the reduced proportion of αSMA- and Notch3-positive cells (Supplemental Figure 5B,C). To confirm Dll4 requirement for coronary arterial formation and circumvent the lethality resulting from E12.5 tamoxifen administration in *Dll4^flox^*;*Pdgfb-iCre^ERT2^*mice, we crossed Dll4^flox^ mice with *Cdh5-Cre^ERT2^* driver line (Wang 2010) to obtain *Dll4^flox^*; *Cdh5-Cre^ERT2^* mice. (Supplemental Figure 5A). Tamoxifen induction at E12.5 resulted in the reduction of coronary arterial coverage, unchanged vein coverage but reduced vein caliber (Supplemental Figure 5D,F). N1ICD staining in the endothelium was decreased by 60% compared with controls (Supplemental Figure 6E,F), likely due to reduced EC contribution to the arteries. Furthermore, *Dll4^flox^*;*Cdh5-Cre^ERT2^* mutant heart coronaries had decreased pervivascular cell coverage as indicated by the reduced proportion of *α*SMA- and Notch3-positive cells (Supplemental Figure 5E,F). The impact of impaired coronary vessel formation in endothelial *Jag1-* or *Dll4-* mutants was also indicated by their diminished ventricular wall thickness (Supplemental Figure 6A).

**Figure 4.**
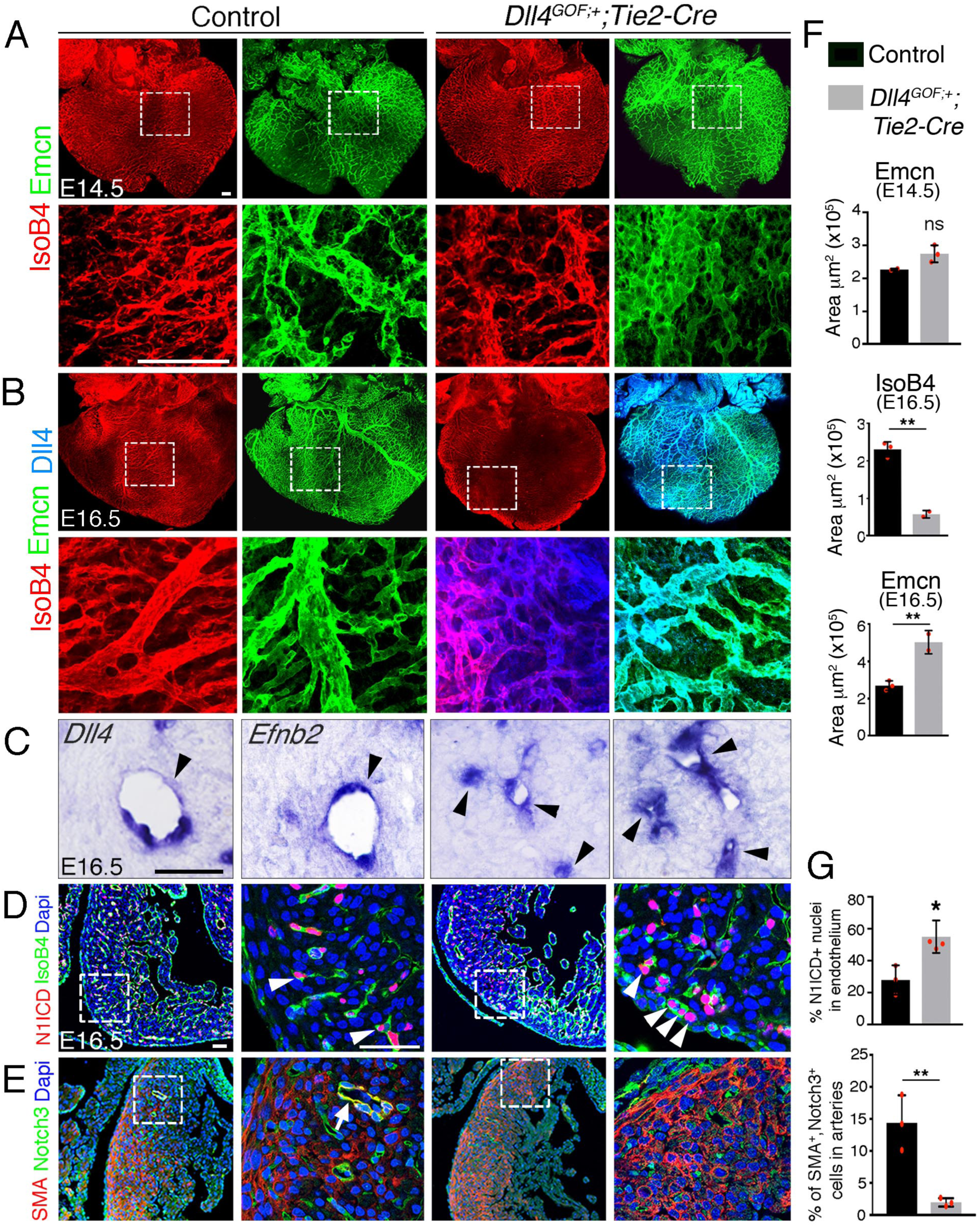
Forced Dll4 expression in endothelium disrupts coronary arteriovenous differentiation and remodeling. **(A)** Dorsal view of whole-mount immunochemistry for Emcn (green) in E11.5 control and *Dll4^GOF^;Tie2-Cre* heart. Scale bar, 100 µm. **(B)** Dorsal view of whole-mount immunochemistry for IsoB4 (red) and Emcn (green) in E14.5 control and *Dll4^GOF^;Tie2-Cre* mutant heart. Scale bar, 100 µm. **(C)** ISH of *Dll4* and *Efnb2* on E16.5 control and *Dll4^GOF^;Tie2-Cre* heart sections. Arrowheads indicate coronary arteries. Scale bar, 50 µm. **(D)** Immunohistochemistry for N1ICD (red) and IsoB4 (green) on E16.5 control and *Dll4^GOF^;Tie2-Cre* heart sections. Dapi counterstain (blue). Arrowheads indicate N1ICD-stained nuclei. Scale bar, 50 µm. **(E)** Immunohistochemistry for SMA (red) and Notch3 (green) on E16.5 control and *Dll4^GOF^;Tie2-Cre* heart sections. Dapi counterstain (blue). The arrow points to a coronary vessel stained by SMA and Notch3. Scale bar, 50 µm. Microscope: Nikon A1-R. Software: NIS Elements AR 4.30.02. Build 1053 LO, 64 bits. Objectives: Plan Apo VC 20x/0,75 DIC N2 dry; Plan Fluor 40x/1,3 Oil DIC H N2 Oil. **(F)** Quantified data for control and *Dll4^GOF^;Tie2-Cre* hearts: E14.5, area covered by veins; E16.5, area covered by coronary arteries, area covered by veins. (**G**) Quantified data of the percentage of N1ICD-positive nuclei in E16.5 endothelium as a percentage (%) of total nuclei; and SMA-Notch3 co-immunostaining in coronary arteries. Data are mean ± s.d, (n= 3 sections from 3 control and n= 4 sections from 3 mutant embryos) \**P* < 0.05, \*\**P* < 0.01, by Student’s *t*-test; ns. not significant.

**Figure 5.**
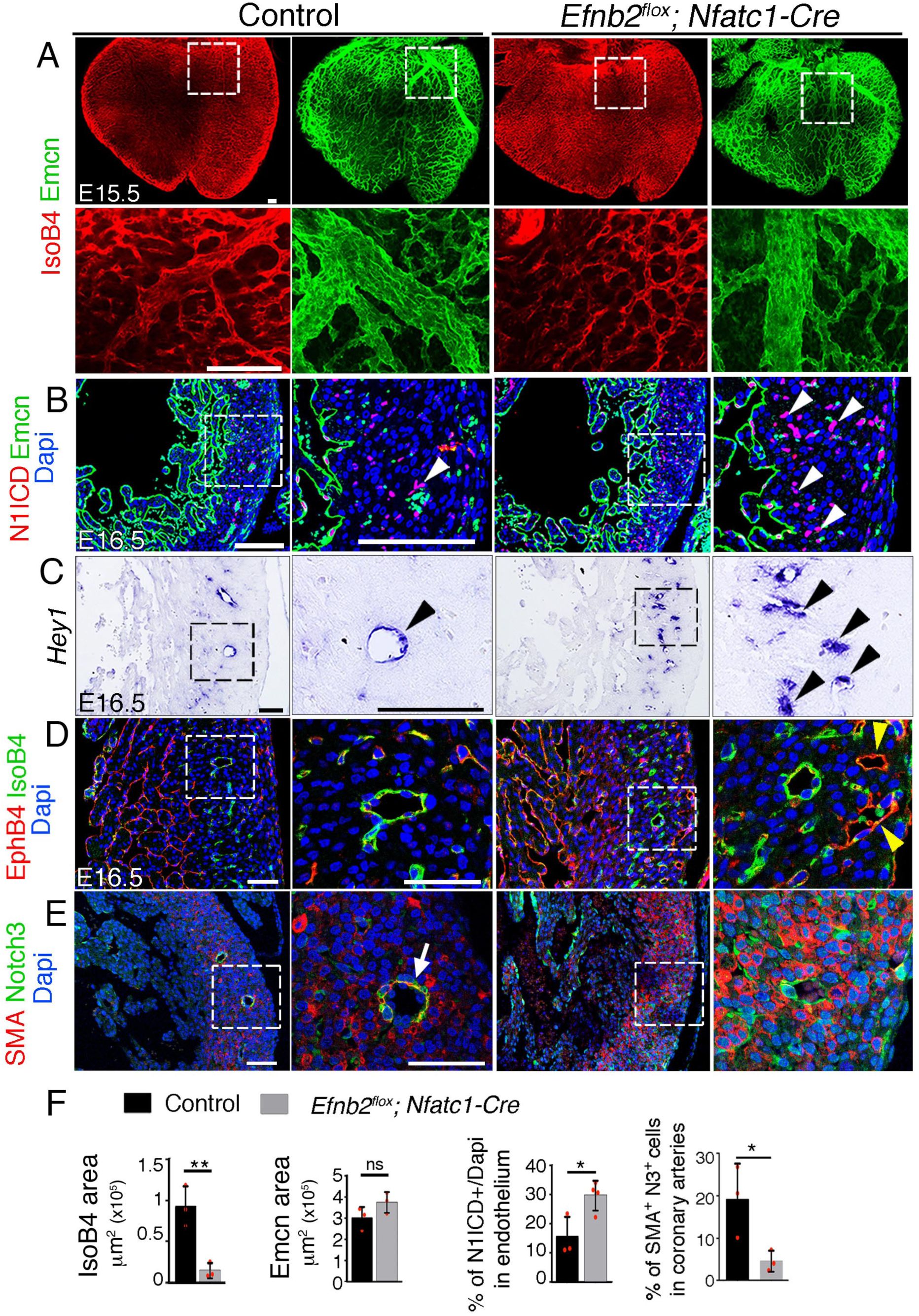
Endocardial *Efnb2* inactivation disrupts coronary artery differentiation and remodeling. **(A)** Dorsal view of whole-mount immunochemistry for IsoB4 (red) and Emcn (green) in E15.5 control and *Efnb2^flox^;Nfatc1-Cre* hearts. Scale bars,100 µm.. **(B)** Immunohistochemistry of N1ICD (red) and Emcn (green) on E16.5 control and *Efnb2^flox^;Nfatc1-Cre* heart sections. Dapi-counterstain (blue). Arrowheads indicate N1ICD-stained nuclei. Scale bars,100 µm. **(C)** ISH of *Hey1* on E16.5 control and *Efnb2^flox^;Nfatc1-Cre* heart sections. Arrowheads indicate *Hey1*-expressing coronaries. Scale bars,100 µm. **(D)** Immunohistochemistry for EphB4 (red) and IsoB4 (green) on E16.5 control and *Efnb2^flox^;Nfatc1-Cre* heart sections. Dapi-counterstain (blue). Yellow arrowheads indicate EphB4-stained vessels. Scale bar, 50 µm. (**E**) Immunohistochemistry for SMA (red) and Notch3 (green) on E16.5 control and *Efnb2^flox^;Nfatc1-Cre* heart sections. Dapi-counterstain (blue). Arrow points to a coronary artery stained by SMA and Notch3. Microscope: Nikon A1-R. Software: NIS Elements AR 4.30.02. Build 1053 LO, 64 bits. Objectives: Plan Apo VC 20x/0,75 DIC N2 dry; Plan Fluor 40x/1,3 Oil DIC H N2 Oil. **(F)** Quantified data for control and E16.5 WT and *Efnb2^flox^;Nfatc1-Cre* hearts: E15.5, area covered by coronary arteries, area covered by veins, and percentage of N1ICD-stained nuclei in endothelium as percentage (%) of total nuclei; E16.5, SMA-Notch3 co-immunostaining in coronary arteries. Data are mean ± s.d. (n= 3 sections from 3 control embryos and n= 3 sections from 3-4 mutant embryos). \**P* < 0.05, \*\**P* < 0.01 by Student’s *t*-test; n.s., not significant.

To examine the effect of endothelial *Dll4* deletion on cardiac gene expression we performed RNA-seq on E15.5 *Dll4^flox^;Cdh5-Cre^ERT2^*mutant ventricles. Therefore of 163 DEGs, 138 were upregulated, while 25 were downregulated (Supplemental Figure 6B; supplemental Table 2, sheet 3). As expected, *Dll4* was downregulated, while *Vegfa* was upregulated, suggesting cardiac hypoxia (Supplemental Figure 6B). *Cdkn1a/p21* was upregulated, indicating impaired cell proliferation (Figure 5B). IPA identified strong upregulation of endothelial cell functions (Supplemental Figure 6B: Supplemental Table 3), whereas analysis of upstream regulators identified an enrichment for transcriptional activators associated with innate inflammatory responses (Irfs and Stats), suggesting defective vascular integrity. Other upstream regulators were associated with cardiovascular disease, including hypertrophy (Nfatc2) and oxidative stress responses (Nfe212; Supplemental Figure 6B right: supplemental Table 3, sheet 3). Moreover, ISH showed reduced endothelial expression of *Efnb2* and *Dll4* in smaller caliber vessels (Supplemental Figure 6C), suggesting a loss of arterial identity, whereas *Fapb4* was upregulated (Supplemental Figure 6C) consistent with increased coronary vessel density of the un-remodeled coronary plexus. Thus, endothelial Jag1 or Dll4 signaling promotes remodeling and maturation of the coronary vascular tree, and in the absence of adequate vascular remodeling, cardiac growth is impaired.

### Forced endothelial *Dll4* expression disrupts coronary vascular remodeling

To further characterize the role of Notch in coronary development, we generated a transgenic line (*Dll4^GOF^*) bearing a *Rosa26-CAG-floxNeoSTOPflox-Dll4-6xMycTag* expression cassette (see *Supplemental Methods*). *Tie2-Cre-*mediated removal of the *floxed NeoSTOP* sequences resulted in a mild 1.2-fold endothelial *Dll4* overexpression that permitted survival of transgenic embryos bearing a single copy of the *Dll4^GOF^*allele (not shown). At E14.5, forced *Dll4* expression lead to marginally increased, sub-epicardial vessel coverage (Figure 4A,F) and vascular malformations similar to those found in *Dll4^flox^*;*Pdgfb-iCre^ERT2^*and *Dll4^flox^;Cdh5-Cre^ERT2^* mutants (Supplemental Figure 3E). However, by E16.5 IsoB4-positive (arteries) intramyocardial vessels were substantially decreased (Figure 4B,F). Accordingly, *Dll4* and *Efnb2* expression was restricted to smaller caliber vessels in *Dll4^GOF^* transgenics (Figure 4C). In contrast, superficial Emcn-positive vessels (veins) were more numerous and dense (Figure 4B,F). N1ICD expression was increased and extended to the prospective veins (Figure 4D,G), consistent with Notch1 gain-of-function in endothelium. Arterial smooth muscle cell coverage was also substantially reduced (Figure 4E,G), suggesting defective pericyte and/or smooth muscle differentiation. These coronary vascular defects were associated with below-normal cardiomyocyte proliferation and reduced myocardial thickness (not shown). Therefore, endothelial *Dll4* overexpression blocks coronary artery formation, vessel remodeling and maturation, and impairs cardiac growth. These results were surprising, as our loss-of-function data support a pro-arteriogenic role for Dll4.

To support our *Dll4^GOF^* results, we used a transgenic line conditionally expressing MFng (*MFng^GOF^*) (D’Amato et al. 2016) to test whether increased MFng activity favoured Dll4-mediated signaling, and thus impaired arteriogenesis. At E11.5, *MFng^GOF^;Tie2-Cre* embryos displayed reduced SV sprouting, increased Notch1 signaling and myocardial wall thinning (Supplemental Figure 7A,B). However, by E14.5 *MFng^GOF^;Tie2-Cre* embryos displayed an extensive vascular network (Supplemental Figure 7C), although the arteries appeared atrophied (Supplemental Figure 7C) and prospective veins appeared more numerous and intricately branched (Supplemental Figure 7C). *MFng^GOF^;Tie2-Cre* embryos displayed numerous capillary malformations at E14.5 (Supplemental Figure 3F), suggesting defective arterial-venous differentiation. The decrease in arterial coverage, and increase in venous coverage in *MFng^GOF^;Tie2-Cre* embryos became more obvious by E16.5 (Supplemental Figure 7C,G). N1ICD expression was markedly increased (Supplemental Figure 7D,G), as expected, while the expression of the Notch targets *HeyL* and *Efnb2* was restricted to smaller caliber vessels (Supplemental Figure 7E). *Vegfa* was upregulated throughout (Supplemental Figure 7E), suggesting that *MFng^GOF^;Tie2-Cre* hearts are hypoxic. Pericyte coverage and smooth muscle differentiation, as shown by *α*SMA and Notch3 co-immunostaining, were significantly decreased at E16.5, (Supplemental Figure 7F,G), suggesting defective pericyte recruitment and/or differentiation. Ink injection confirmed that *MFng^GOF^;Tie2-Cre* coronaries were “leaky” (Supplemental Figure 7H), and therefore functionally deficient.

### EphrinB2 is required for coronary arteriogenesis and vessel branching

*Efnb2* is necessary for arterial-venous differentiation and vascular maturation (Kania and Klein 2016) and is a Notch target during ventricular chamber development (Grego-Bessa et al. 2007). We find *Efnb2* expression in ventricular endocardium at E10.5 (Supplemental Figure 8A) but not in SV endothelium (not shown). At E13.5, *Efnb2* was found in emerging arteries and prospective veins (Supplemental Figure 8A), and at E16.5 was confined to arteries (Supplemental Figure 8A). Therefore, *Efnb2* is expressed dynamically in a pattern similar to *Dll4* and *MFng*.

To examine EphrinB2 function in coronary angiogenesis, we crossed mice bearing a conditional *Efnb2^flox^* allele with the *Nfatc1-Cre* line. Whole-mount IsoB4 immunostaining revealed reduced artery coverage in E15.5 *Efnb2^flox^;Nfatc1-Cre* hearts (Figure 5A,F), while vein coverage was unchanged (Figure 5A,F). Histological examination showed that the compact myocardium in the ventricles was 30% thinner at E16.5 (Supplemental Figure 8B,C), which could be attributed to the reduced proliferation observed at E14.5 (Supplemental Figure 8C,D). The presence of smaller caliber arteries coincided with higher Notch1 activity (Figure 5B,F) and increased *Hey1* expression (Figure 5C), suggesting negative feedback between EphrinB2 and Notch signaling. Although the patterning of prospective veins was unchanged we found occasional malformations (Supplemental Figure 3G,H) and transmural communications or shunts between the endocardium and epicardium, which we interpret to be arteriovenous fistulae (Supplemental Figure 8F). Accordingly, the venous marker EphB4 was expressed ectopically in a subset of intramyocardial vessels at E16.5 (Figure 5D; supplementary Figure 8E), indicative of abnormal venous identity. *Vegfa* was upregulated throughout the heart (Supplemental Figure 8F), suggesting that *Efnb2^flox^;Nfatc1-Cre* hearts are hypoxic. Arterial smooth muscle coverage was reduced substantially (Figure 5E,F), suggesting defective vessel maturation. Die perfusion in *Efnb2^flox^;Nfatc1-Cre* heart*s* revealed absence of left anterior descending coronary artery at E16.5 (Supplemental Figure 8G) and ink perfusion revealed vascular hemorrhaging in the most distal section of the coronary arterial tree at E18.5 (Supplemental Figure 8H), consistent with vessel leakiness, while aortic connections were normal (Supplemental Figure 8H). Thus, endocardial EphrinB2 is required for coronary arterial remodeling, vessel structural integrity, and cardiac growth.

### EphrinB2 mediates Jag1 and Dll4 signaling in arterial branching morphogenesis

To model coronary angiogenesis *ex vivo*, we developed a ventricular explant assay (Figure 6A; (Zhang and Zhou 2013). Compared with controls, *Jag1^flox^;Nfatc1-Cre* explants had a lower angiogenic potential, manifested as smaller caliber vessels, longer distances between branching points and a lower number of transmural endothelial branches (Figure 6B,C,K). Conversely, *Dll4^flox^;Nfatc1-Cre* or *Notch1^flox^;Nfatc1-Cre* explants yielded more densely interconnected vascular networks of larger vessel caliber, longer distances between vessel branching points, and more transmural endothelial branches (Figure 6B,D,E,K). *Notch1^flox^;Nfatc1-Cre* explants displayed a phenotype similar to *Dll4^flox^;Nfatc1-Cre* explants, although the differences in vessel caliber and endothelial branch number did not reach significance (Figure 6B,E,K). *MFng^GOF^;Tie2-Cre* and *Dll4^GOF^;Tie2-Cre* explants showed a broadly similar phenotype, with substantial lengthening of branching point distance and a below-normal number of endothelial branches (Figure 6B,I,J,K). In contrast, *MFng^GOF^;Tie2-Cre* explants had smaller caliber vessels than *Dll4^GOF^;Tie2-Cre* explants (Figure 6I,J,K). *Efnb2^flox^;Nfatc1-Cre* explants formed sparsely interconnected networks with decreased vessel caliber, reduced number of transmural endothelial branches, and increased distances between branching points, similar to those seen in *Jag1^flox^;Nfatc1-Cre* explants (Figure 6B,F,K).

**Figure 6.**
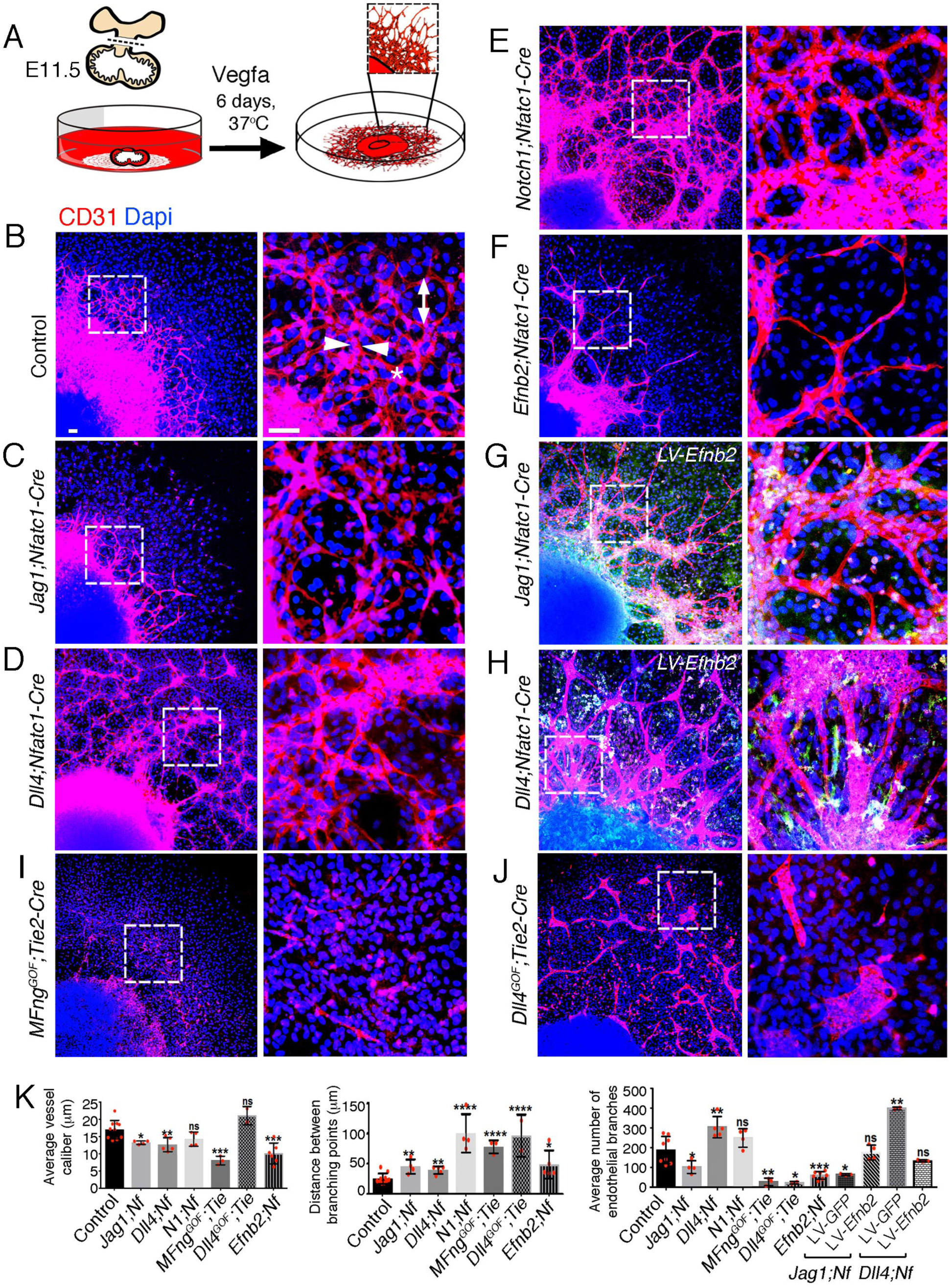
EPHRINB2 rescues disrupted arterial branching in endocardial explants from *Jag1* and *Dll4* mutant hearts. **(A)** Ventricular explant assay procedure. **(B-J)** Representative images of E11.5 cultured ventricular explants from the following embryos: **(B)** control (n=8-10), **(C)** *Jag1^flox^;Nfatc1-Cre* (n=4), **(D)** *Dll4^flox^;Nfatc1-Cre* (n=5), **(E)** *Notch1^flox^;Nfatc1-Cre* (n=4), **(F)** *MFng^GOF^;Tie2-Cre* (n=3), **(G)** *Dll4^GOF^;Tie2-Cre* (n=3), (**H**) *Efnb2^flox^;Nfatc1-Cre* (n=5-7), **(I)** *Dll4^flox^;Nfatc1-Cre* infected with *Efnb2-* overexpressing lentivirus (n=5-7). Microscope: Nikon A1-R. Software: NIS Elements AR 4.30.02. Build 1053 LO, 64 bits. Objectives: Plan Apo VC 20x/0,75 DIC N2 dry; Plan Fluor 40x/1,3 Oil DIC H N2 Oil. **(J)** Quantification of vessel caliber, branching point distance, and mean endothelial branch number. Data are means ± s.d. \*\*\*\**P* < 0.0001; \*\*\**P* < 0.001; \*\**P* < 0.01; \**P* < 0.05, by Student’s t-test; n.s not significant. Scale bars, 100 µm.

Based on the congruence of endocardial/endothelial *Jag1*, *Dll4* and *Efnb2* loss-of-function phenotypes, we tested whether EphrinB2 mediates Notch function in coronary angiogenesis. Thus, delivery of a lentivirus expressing *Efnb2* to *Jag1^flox^;Nfatc1-Cre* ventricular explants normalized the reduced endothelial branch number to the number seen in control explants (Fig. 6B,G,K). Likewise, *Efnb2* expression restored the elevated number of endothelial branches in *Dll4^flox^;Nfatc1-Cre* explants to control levels (Fig. 6B,H,K), showing that the branching defect could be normalized in both mutants.

### EphrinB2 functions downstream of Jag1 and Dll4 in capillary tube formation

The ventricular explant assay assesses the collective behaviors of endocardium and coronary endothelial outgrowth. In order to determine endothelial behavior exclusively, we performed capillary tube formation assays with primary human endothelial cells (HUVEC). shRNA directed against JAG1, DLL4 or EFNB2 were lentivirally-transduced into HUVEC. This resulted in reduction of mRNA levels of 30% and 60% for JAG1 and DLL4 respectively (Figure 7A), and of 50% for EFNB2 (Supplemental Figure 9A). The NOTCH target HEY1 was unchanged in *shJAG1*-infected cells and 50% decreased in *shDLL4*-infected cells (Figure 7A). The activity of EFNB2 lentivirus was verified by rescuing shRNA-mediated EFNB2 knockdown (Supplemental Figure 9B). EFNB2 knockdown resulted in decreased capillary network complexity (Supplemental Figure 9B,C). These parameters were restored to control levels by lentiviral-mediated EFNB2 overexpression (Supplemental Figure 9B,C).

**Figure 7.**
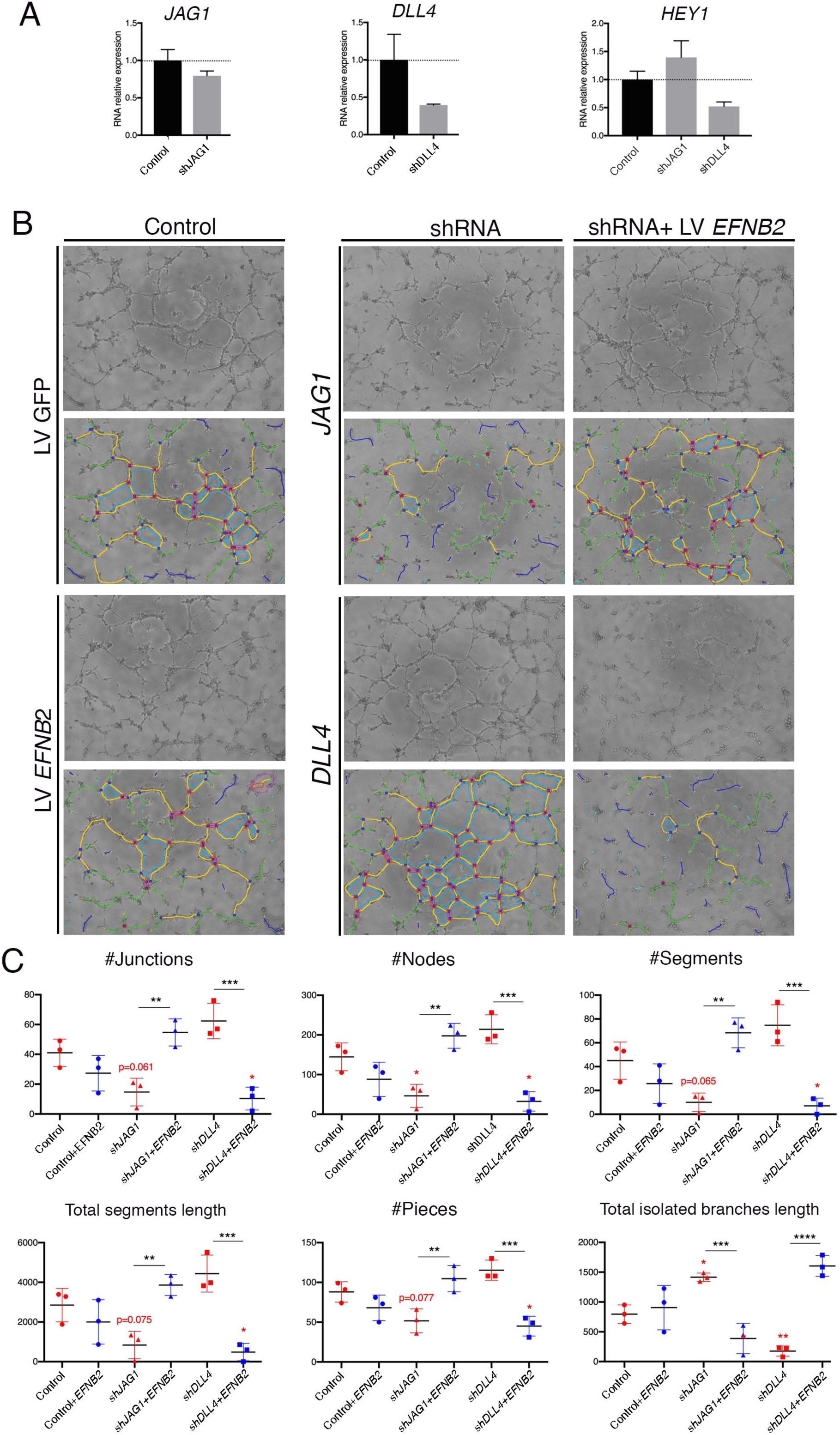
EPHRINB2 rescues defective capillary network formation resulting from shRNA-mediated silencing of JAG1 and DLL4 in HUVEC. **(A)** qRT-PCR of JAG1, DLL4 and HEY1 after transduction of shRNA. **(B)** Representative phase contrast images (1 of 2 experiments) of the HUVEC network, after transduction of indicated shRNA and rescue by EFNB2 analyzed by the Angiogenesis Analyzer for ImageJ. **(C)** Significantly changed measurements in the analyzed area: nodes surrounded by junctions (red surrounded by blue). Isolated elements (purple). Segments (yellow). Number of branches (green) and meshes (light blue) were not significantly altered (not shown). Total segments length: sum of length of the segments. Number of pieces: sum of number of segments, isolated elements and branches detected. Total isolated branches length: sum of the length of the isolated elements. Red asterisks refer to comparisons between experimental and control situations. Black asterisks refer to comparisons between shRNA-mediated inhibition and LV-mediated rescue. Data are means ± s.d. \*\*\*\**P* < 0.0001; \*\*\**P* < 0.001; \*\**P* < 0.01; \**P* < 0.05, by ANOVA.

We next tested whether EPHRINB2 acts downstream of JAG1 and DLL4 in capillary tube formation. JAG1 knockdown inhibited capillary tube formation and decreased network complexity as determined by measuring EC junctions, nodes, segments and pieces that were below control, while the “total isolated branches length” was above control (Figure 7B,C). However, these parameters were restored to control levels in presence of the EFNB2 transgene (Figure 7B,C), suggesting that EPHRINB2 compensates for JAG1 knockdown in this assay. In contrast, knockdown of DLL4 increased capillary tube formation and network complexity (junctions, nodes, segments and pieces; Figure 7B,C), while measurements were restored and even went beyond control by overexpressing EFNB2 (Figure 7B,C). Thus, EFNB2 overexpression not only compensates for the absence of DLL4 in this assay, but has an added effect as well.

Taken together, our observations with ventricular explants and HUVEC indicate that JAG1 and EPHRINB2 promote coronary vessel sprouting and branching, whereas DLL4, MFNG and NOTCH1 inhibit these processes. Moreover, EPHRINB2 mediates signaling from both JAG1 and DLL4 to regulate endothelial branching during coronary angiogenesis.

## Discussion

The origin of the coronary endothelium from a venous source has prompted the suggestion that arteries need to be “reprogrammed” from veins (Red-Horse et al. 2010). We found Jag1, Dll4, MFng, and N1ICD expression in SV endothelium and sub-epicardial capillaries, with Dll4 and MFng expression maintained subsequently in prospective veins. These Notch pathway elements are co-expressed with the venous marker endomucin, which is also expressed in the endocardium and SV (Cavallero et al. 2015). This implies that nascent subepicardial vessels have a mixed arterial/venous identity. During embryonic angiogenesis, Notch is necessary for artery-vein specification—the reversible commitment of ECs to arterial or venous fate—before the onset of blood flow (Swift and Weinstein 2009). Our finding of SV endothelial cell heterogeneity in relation to arterial-venous identity is consistent with the notion that early vascular beds are phenotypically plastic during embryonic development (Moyon et al. 2001; Chong et al. 2011; Fish and Wythe 2015). Our results are consistent with recently published data showing that pre-arterial cells expressing Notch pathway elements are present in coronary endothelium (Su et al. 2018), prior to blood flow onset.

We show that Notch ligands Jag1 and Dll4 are required for SV sprouting angiogenesis, further lending support to the notion that coronary arteries are specified by Notch prior to blood flow (Su et al. 2018). Dll4 is the key Notch ligand activator regulating embryonic (Duarte et al. 2004; Gale et al. 2004; Krebs et al. 2004; Benedito et al. 2008) and postnatal retinal angiogenesis (Hellstrom et al. 2007; Lobov et al. 2007), whereas Jag1 antagonizes Dll4-Notch1 signaling (Benedito et al. 2009) and acts downstream of Dll4-Notch1 to promote smooth muscle differentiation (Pedrosa et al. 2015). *Jag1* or *Dll4* inactivation in the SV results in arrested and excessive angiogenesis, respectively, resembling the situation in the retina (Benedito et al. 2009). Moreover capillary “entanglements” suggestive of vascular malformations were present in the early plexus of *Jag1^flox^;Nfatc1-Cre* and *Dll4^flox^;Nfatc1-Cre* mutants. Vascular malformations in Notch mutants have been attributed to unresolved intermingling of arteries and veins during differentiation, or to failed maintenance of arterial and venous identities in the vascular bed (Gale et al. 2004; Krebs et al. 2004). The presence of these malformations in *Jag1^flox^;Nfatc1-Cre* and *Dll4^flox^;Nfatc1-Cre* mutants could therefore indicate that endothelial progenitors begin to differentiate into arteries and veins very soon after exiting the SV.

Consistent with the known roles of the Notch ligands in angiogenesis, the hierarchical organization of the coronary vascular tree was profoundly altered in the late-induced *Jag1* and *Dll4* mutants, implying that both Notch ligands are required for high-order complexity of the coronary vessels. *Jag1* or *Dll4* inactivation during coronary arterial remodeling results in smaller-diameter arteries, consistent with Notch promoting vascular remodeling and arterial fate commitment, as described in other developmental settings (Duarte et al. 2004; Gale et al. 2004; Krebs et al. 2004). However, gain of Notch function also led to smaller caliber arteries, when the opposite outcome was anticipated (Uyttendaele et al. 2001; Trindade et al. 2008; Krebs et al. 2010). This result implies that non-physiological variations of Notch activity lead to the arrest of coronary artery differentiation. *Dll4* mutants also showed coronary vessel hemorrhaging reminiscent of that found in vascular disorders (Park-Windhol and D’Amore 2016) or tumors (Goel et al. 2011) and indicative of disrupted endothelial integrity.

We reasoned that the coronary maturation defects observed in *Jag1* or *Dll4* mutants are due, at least in part, to the loss of EphrinB2 function given that *Efnb2* is a direct Dll4-Notch signaling target in ECs (Iso et al. 2006) and endocardium during ventricular development (Grego-Bessa et al. 2007). Moreover, EphrinB2 and VEGF are functionally linked during angio- and lymphangiogenesis; EphrinB2 is a direct activator of VEGFR2 and VEGFR3, and therefore cooperates in the mechanism leading to tip cell extension and vessel sprouting (Sawamiphak et al. 2010; Wang et al. 2010). Thus, *Efnb2* inactivation leads to coronary artery remodeling defects, similar to those resulting from *Jag1* or *Dll4* inactivation, suggesting that EphrinB2 functions downstream of Notch to promote coronary arterial remodeling. This notion is supported by the angiogenic ventricular explant and EC capillary tube assays, in which opposite effects of *Jag1* or *Dll4* deficiency on vessel branching are rescued by transduction of an EFNB2-expressing lentivirus, identifying EphrinB2 as a Notch effector during coronary artery development.

A common feature among the endothelial Notch loss- and gain-of-function models analyzed in our study is the thin ventricular wall, associated with defective compact myocardium proliferation. Of interest is that myocardial inactivation of *Jag1*, or combined inactivation of *Jag1* and *Jag2*, or *Mib1*, leads to thinner ventricular walls, accompanied by reduced cardiomyocyte proliferation, disrupted ventricular chamber patterning, and cardiomyopathy (Luxan et al. 2013; D’Amato et al. 2016). In contrast, chamber patterning is maintained in endocardial *Jag1*, *Dll4*, and *Efnb2* mutants and endothelial *Dll4* (D’Amato et al. 2016) and *Jag1* mutants, indicating that reduced ventricular wall thickness in these mutants is a consequence of the lack of a well-developed coronary system. These observations suggest that coronaries provide metabolic substrates or “angiocrine signals” (Rafii et al. 2016) for compact myocardium growth, rather than patterning signals. The “trophic” role of newly formed coronaries may be required for ventricular growth before perfusion from the aorta, and lack of adequate trophic support may explain the early lethal phenotype of *Jag1^flox^*- and *Dll4^flox^;Nfactc1-Cre* mutants. In addition, our data suggest that ventricular compaction relies on two interconnected Notch-dependent processes: patterning and maturation of the ventricles, and timely development of a functional coronary vessel network, as previously suggested (D’Amato et al. 2016). This may be clinically relevant to the study and treatment of cardiomyopathies.

## Acknowledgments

We thank the CNIC Genomics Unit for RNA-seq experiments and S. Bartlett (CNIC) for English editing. This study was supported by grants SAF2016-78370-R, CB16/11/00399 (CIBER CV), and RD16/0011/0021 (TERCEL) from the Spanish Ministry of Science, Innovation and Universities (MCIU) and grants from the Fundación BBVA (Ref.: BIO14_298) and Fundación La Marató (Ref.: 20153431) to JLDLP. The cost of this publication was supported in part with funds from the ERDF. The CNIC is supported by the MCIU and the Pro-CNIC Foundation and is a Severo Ochoa Center of Excellence (SEV-2015-0505).

## Authorship contributions

S.I.T.,V.L.O. and V.B. performed experiments, and S.I.T. generated figures. B.P. generated the *Dll4^GOF^* mice. M.J.G. and F.S.C. analyzed the RNA-seq data and generated plots. D.M. and J.L.d.l.P. designed experiments, reviewed the data, and wrote the manuscript. All authors reviewed the manuscript.

## Conflict of Interest Disclosures

The authors declare no competing financial interests.

## Supplemental Material

**Supplemental Figure 1.**
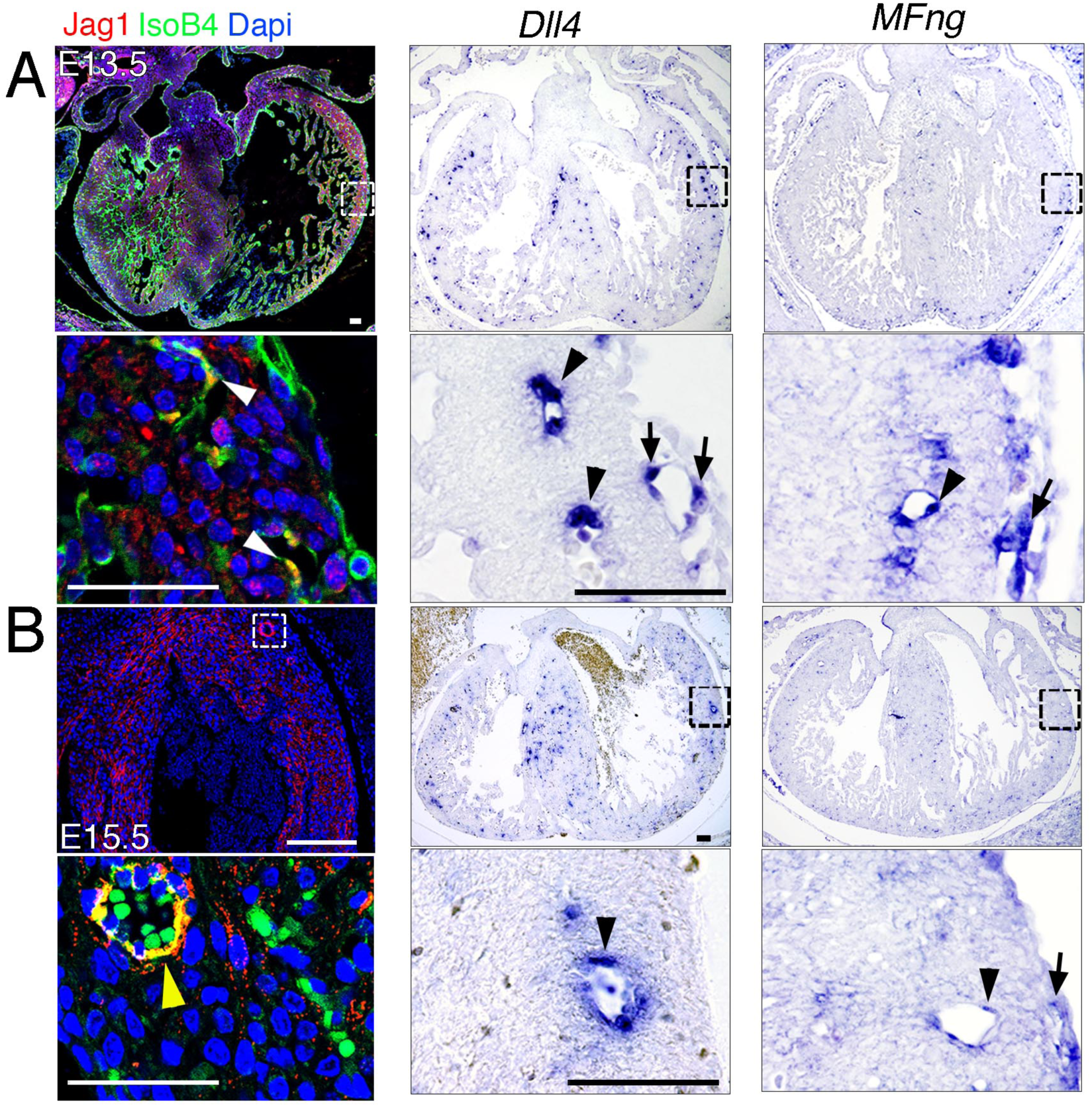
J*a*g1, *Dll4*, and *MFng* expression in developing coronary vessels. **(A)** E13.5 control heart. *Left,* immunohistochemistry for Jag1 (red) and IsoB4 (green); the magnified view shows co-staining of Jag1 and IsoB4 in coronary arteries (white arrowheads). *Center* and *right*, ISH for *Dll4* and *MFng*; magnified views show expression in coronary arteries (black arrowheads) and sub-epicardial vein (arrows). **(B)** E15.5 control heart. Immunohistochemistry for Jag1 (red; top) and IsoB4 co-staining (green; bottom); the magnified view shows co-staining of Jag1 and IsoB4 in a coronary artery (yellow arrowhead). Dapi counterstain (blue). ISH for *Dll4* and *MFng* expression; magnified views show expression in a coronary artery (black arrowhead) and sub-epicardial vein (arrow). Scale bars, 50 µm. Images for Jag1 immunstainings were acquired on Nikon A1-R microscope. Software: NIS Elements AR 4.30.02. Build 1053 LO, 64 bits. Objectives: Plan Apo VC 20x/0,75 DIC N2 dry; Plan Fluor 40x/1,3 Oil DIC H N2 Oil.

**Supplemental Figure 2.**
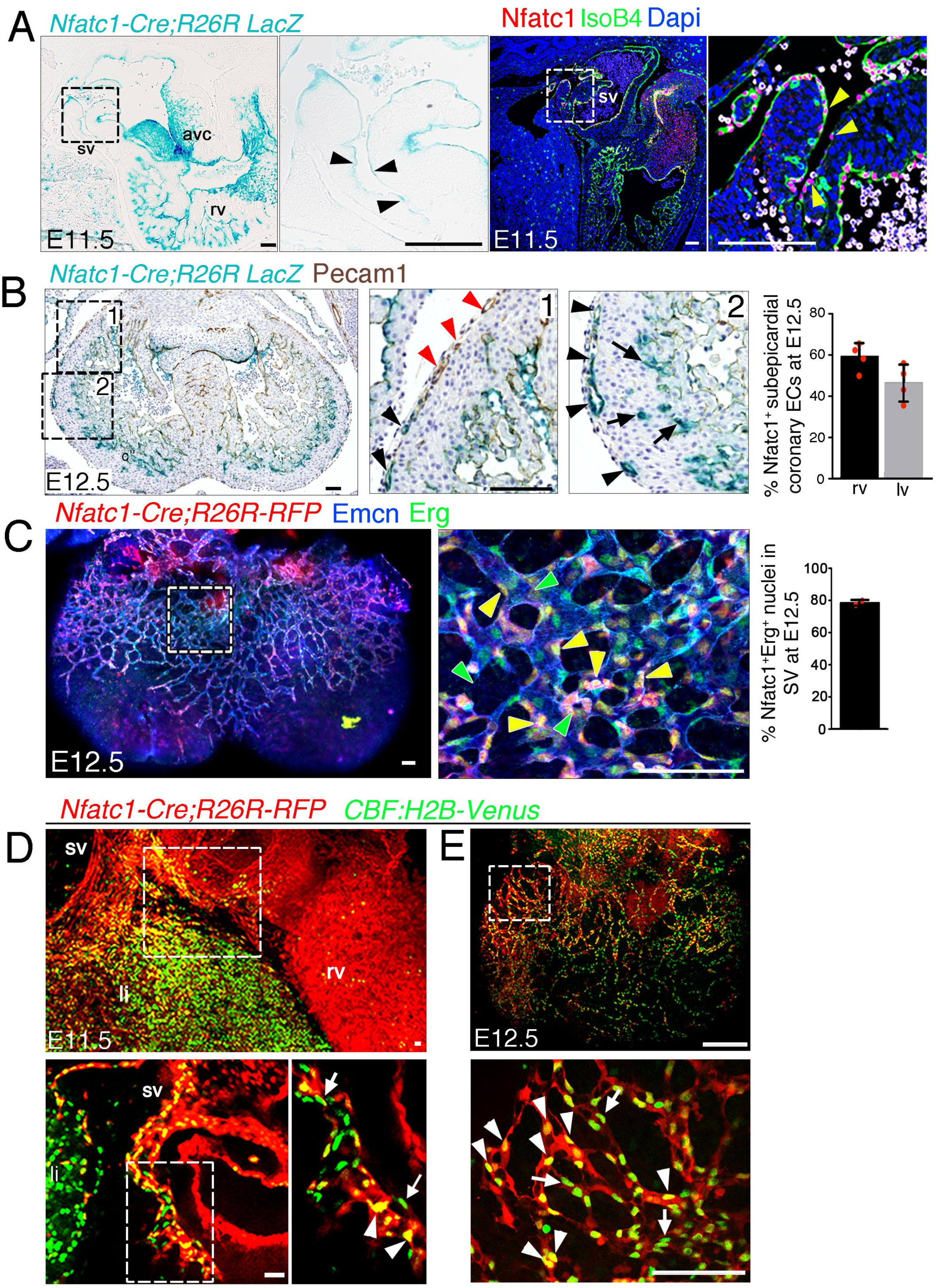
Most ventricular free-wall coronary vessels derive from Nfatc1^+^ progenitors Nfatc1^+^ progenitors give rise to most coronary vessels. **(A)** *Left:* Sagittal section of a E11.5 *Nfatc1-Cre;R26R-LacZ* heart, showing LacZ expression in the sinus venosus (sv; arrowheads in inset), right ventricle (rv), and atrioventricular canal (avc). *Right:* Sagittal section of a E11.5 control heart immunostained for Nfatc1 (red) and IsoB4 (green); yellow arrowheads indicate nuclear Nfatc1 staining in SV endothelial cells. Dapi counterstain (blue). Scale bars 100 µm**. (B)** E12.5 *Nfatc1-Cre;R26R-LacZ* heart co-stained for LacZ and Pecam-1. (1) LacZ staining is weak in vessels proximal to the AVC (red arrowheads) and strong in distal vessels (1, 2, black arrowheads). (2) LacZ staining in vessels close to chamber endocardium is consistent with an endocardial origin of coronary arteries (arrows). The chart shows Nfatc1-positive vessels (*Nfatc1-Cre*-driven LacZ expression) as a percentage of all sub-epicardial vessels (Pecam-1-stained). Data are mean +/-s.d. (n=3 sections from 4 embryos). **(C)** Whole-mount E12.5 *Nfatc1-Cre;Rosa26-RFP* heart (dorsal view), showing RFP fluorescence (red) and immunhistochemistry for Emcn (blue) and Erg (green). Sub-epicardial vessel endothelium contains *Nfatc1-Cre;Rosa26-RFP*-positive nuclei (yellow arrowheads) and *Nfatc1-Cre;Rosa26-RFP-*negative nuclei (green arrowheads). Dapi counterstain (blue). The chart shows Nfatc1-Erg dual-positive nuclei as a percentage of all nuclei in subepicardial vessels. Data are mean ± s.d. (n= 2 hearts). Scale bars, 100 µm. **(D)** Whole-mount E11.5 *Nfatc1-Cre;Rosa26-RFP*:*CBF:H2B-Venus* heart (dorsal view), showing RFP reporter fluorescence (red) and immunohistochemistry for GFP (green). The Z-stack of SV endothelium shows *Nfatc1-Cre;Rosa26-RFP*-*CBF:H2B-Venus* double-positive nuclei (arrowheads) and *CBF:H2B-Venus* single-positive nuclei (arrows). sv, sinus venosus; rv, right ventricle; li, liver. **(E)** Whole-mount E12.5 *Nfatc1-Cre;Rosa26-RFP*-*CBF:H2B-Venus* heart (dorsal view), showing RFP reporter expression (red) and immunohistochemistry for GFP (green). The sub-epicardial vessel endothelium contains *Nfatc1-Cre;Rosa26-RFP*, *CBF:H2B-Venus* dual-positive nuclei (white arrowheads) and *CBF:H2B-Venus* single positive nuclei (arrows). Scale bars, 100 µm. Images for fluorescence immunostainings were acquired on a Nikon A1-R microscope. Software: NIS Elements AR 4.30.02. Build 1053 LO, 64 bits. Objectives: Plan Apo VC 20x/0,75 DIC N2 dry; Plan Fluor 40x/1,3 Oil DIC H N2 Oil.

**Supplemental Figure 3.**
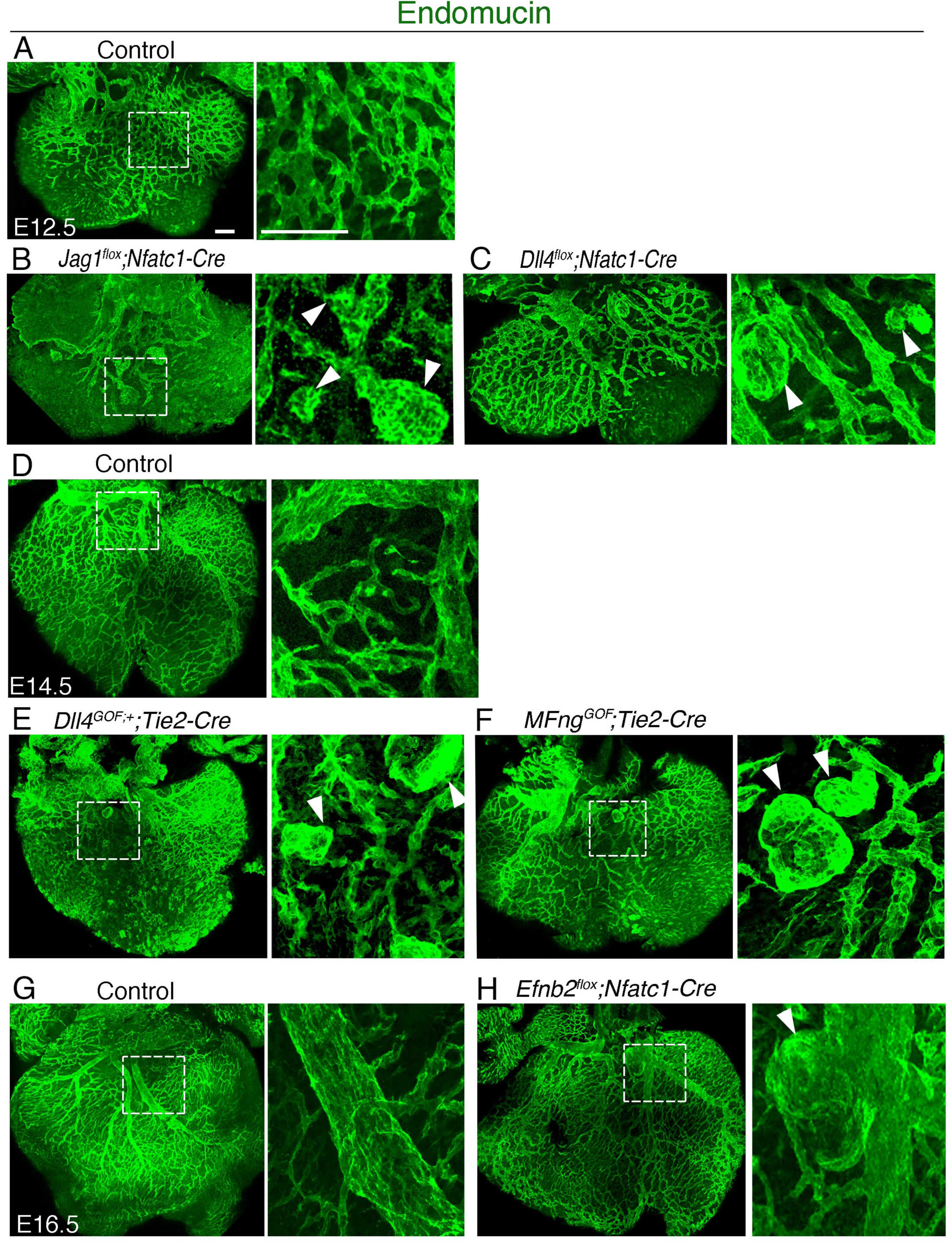
Coronary vessels of Notch pathway and Notch effector mutants display vascular malformations. Dorsal views of whole-mount immunostaining for Emcn (green). **(A)** E12.5 control heart, **(B)** E12.5 *Jag1^flox^;Nfatc1-Cre* heart, **(C)** E12.5 *Dll4^flox^;Nfatc1-Cre* heart, **(D)** E14.5 control heart, **(E)** E14.5 *MFng^GOF^;Tie2-Cre* heart, **(F)** E14.5 *Dll4^GOF^;Tie2-Cre* heart, **(G)** E16.5 control heart, **(H)** E16.5 *Efnb2^flox^;Nfatc1-Cre* heart. Scale bars, 100 µm.

**Supplemental Figure 4.**
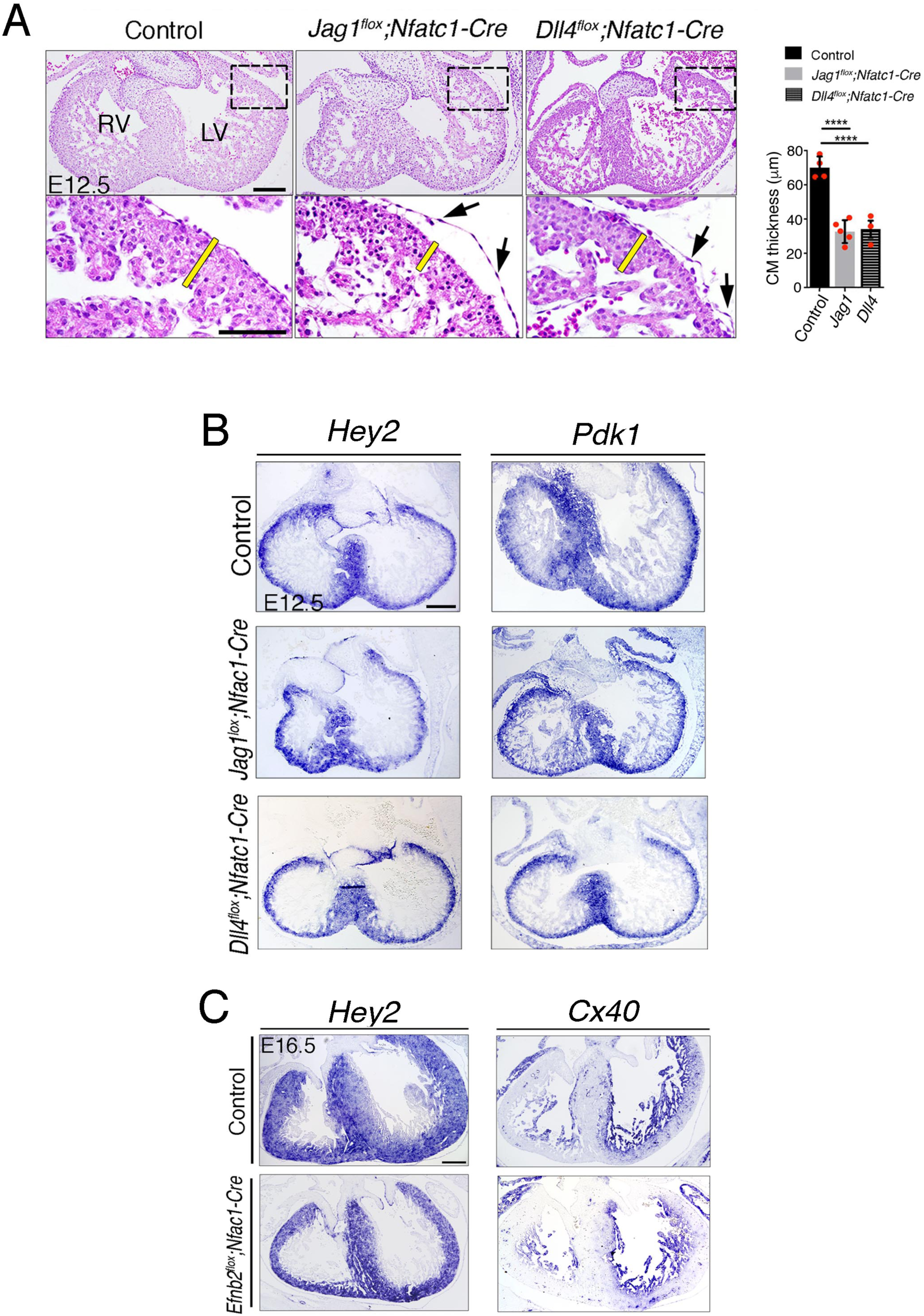
**(A)** Haematoxylin–eosin (H&E) staining on E12.5 control, *Jag1^flox^;Nfatc1-Cre*, and *Dll4^flox^;Nfatc1-Cre* heart sections. Arrows point to detached epicardium. Yellow bars indicate compact myocardium (CM) thickness. Quantified data of compact myocardium (CM) thickness in E12.5 control, *Jag1^flox^;Nfatc1-Cre* and *Dll4^flox^;Nfatc1-Cre* hearts. Data are mean ± s.d. (n=4 control hearts and n=3-5 mutant hearts). ****P < 0.0001 by Student’s *t*-test. (B) Endocardial Jag1 or Dll4 deletion does not affect myocardial patterning. ISH of the compact myocardium markers *Hey2* and *Pdk1* on E12.5 control, *Jag1^flox^;Nfat-Cre*, and *Dll4^flox^;Nfatc1-Cre* heart sections **(C)** ISH of the compact myocardium marker *Hey2,* and trabecular myocardium marker *Cx40* on E16.5 control and *Efnb2^flox^;Nfatc1-Cre* heart sections. Scale bars, 100 µm.

**Supplemental Figure 5.**
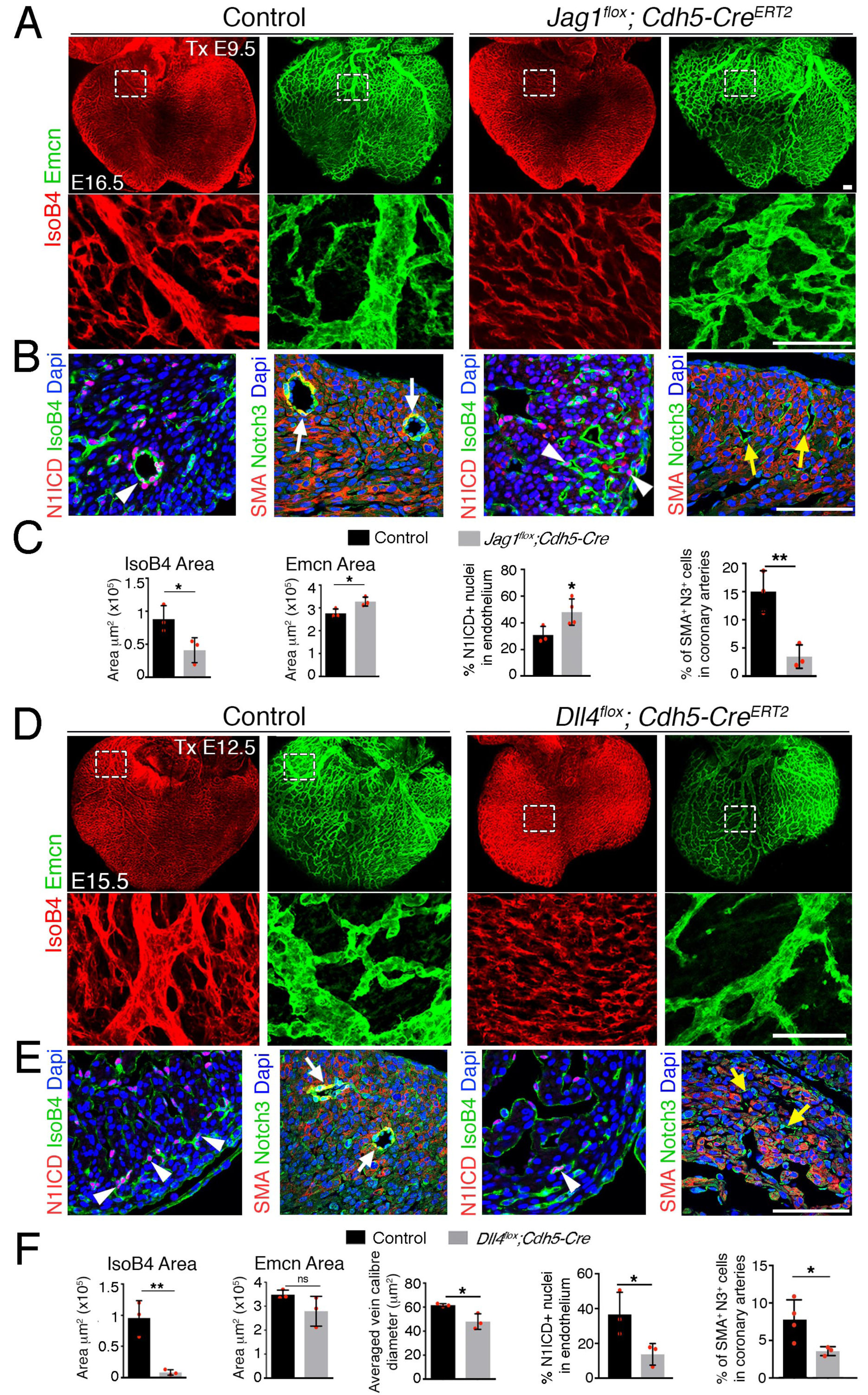
Late endothelial *Jag1* or *Dll4* inactivation disrupts coronary plexus remodelling. **(A)** Dorsal view of whole-mount immunochemistry for Emcn (green) and IsoB4 (red) in E16.5 control and *Jag1^flox^;Cdh5-Cre^ERT2^*mutant hearts, tamoxifen (Tx)-induced at E9.5. Scale bars, 100 µm. **(B)** E16.5 control and *Jag1^flox^;Cdh5-Cre^ERT2^* mutant heart sections. Left, Immunostaining for N1ICD (red) and IsoB4 (green). Right, α-smooth-muscle actin (SMA, red) and Notch3 (green). Dapi counterstain (blue). Arrowheads point to N1ICD-positive nuclei. Arrows point to SMA-Notch3 co-immunostaining. Yellow arrows point to small calibre intramyocardial vessels in Jag1 mutant. **(C)** Quantified data from E15.5 control and *Jag1flox;Cdh5-CreERT2* hearts: area of coverage by coronary arteries and veins; N1ICD-positive nuclei as a percentage of total endothelial nuclei; and SMA-Notch3 co-immunostaining in coronary arteries. **(D)** Whole-mount dorsal view of immunohistochemistry for Emcn (green) and IsoB4 (red) in E15.5 control and *Dll4^flox^;Cdh5-Cre^ERT2^* mutant hearts Tx-induced at E12.5. **(E)** Immunohistochemistry on the left for N1ICD (red) and IsoB4 (green), and on the right for SMA (red) and Notch3 (green) on control E15.5 WT and *Dll4^flox^;Cdh5-Cre^ERT2^*mutant heart sections. Dapi counterstain (blue). Arrowheads indicate N1ICD-positive nuclei. Arrows point to SMA-Notch3 co-immunostaining. Yellow arrows point to small calibre intramyocardial vessels in the Dll4 mutant. Scale bars as in **(A). (F)** Quantified data from E16.5 control and *Dll4^flox^;Cdh5-Cre^ERT2^* hearts: area covered by coronary arteries; area covered by veins; mean vein caliber; percentage of N1ICD-positive nuclei in endothelium as a percentage (%) of total nuclei; and SMA-Notch3 co-immunostaining in coronary arteries. Data are mean +/-s.d. (n= 3 sections from 3 Control embryos and from 3-4 mutant embryos). *P < 0.05, **P < 0.01, by Student’s t-test; n.s., not significant. Scale bars, 100 µm. Microscope: Nikon A1-R. Software: NIS Elements AR 4.30.02. Build 1053 LO, 64 bits. Objectives: Plan Apo VC 20x/0,75 DIC N2 dry; Plan Fluor 40x/1,3 Oil DIC H N2 Oil.

**Supplemental Figure 6.**
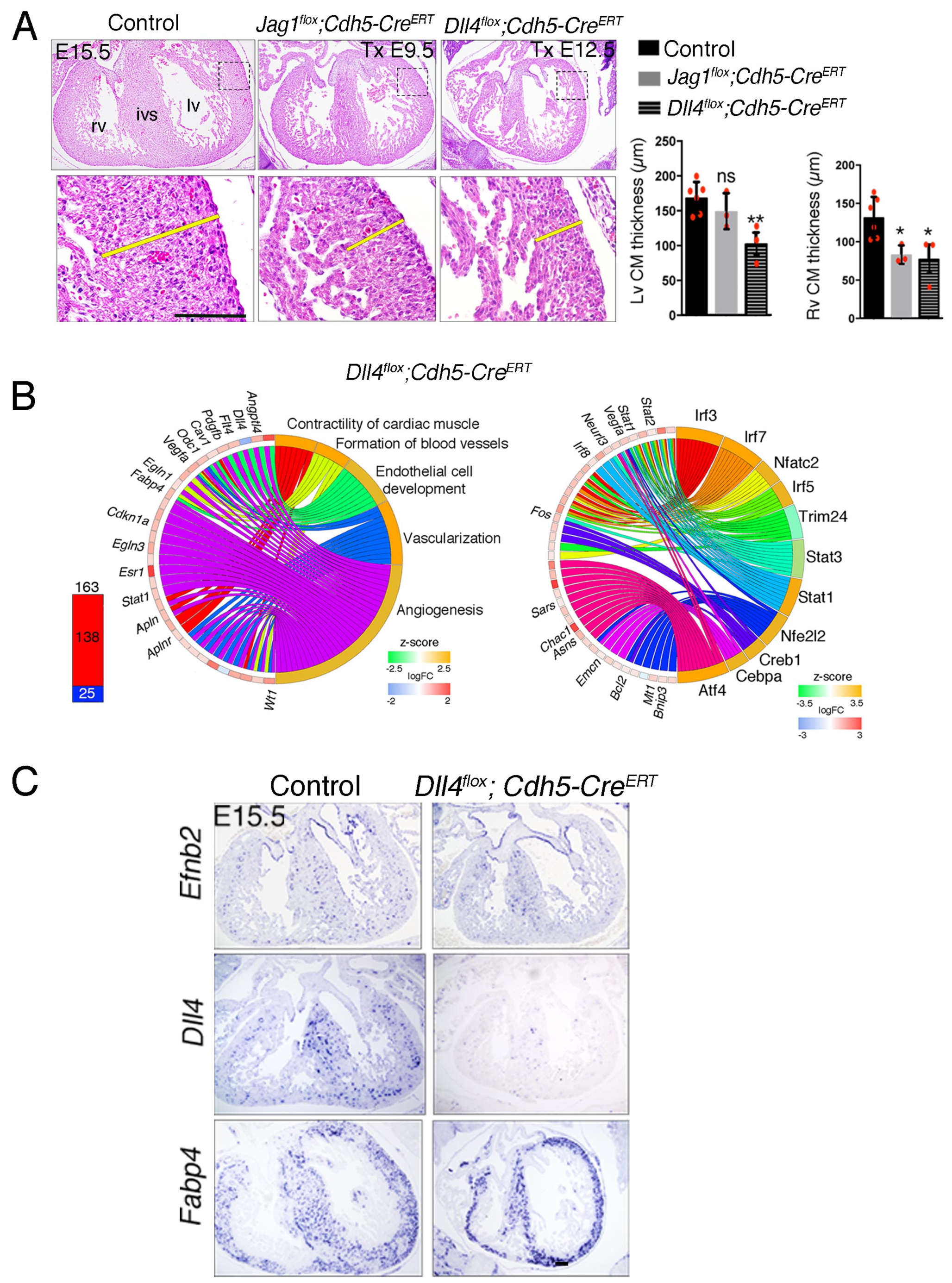
Induced endothelial *Jag1* or *Dll4* deletion disrupts myocardial growth. **(A)** H&E staining of transverse sections of E15.5 hearts from control mice, *Jag1^flox^;Cdh5-Cre^ERT2^*mice tamoxifen (Tx)-induced at E9.5 and *Dll4^flox^;Cdh5-Cre^ERT^* mice Tx-induced at E12.5. Yellow bars indicate compact myocardium (CM) thickness. Charts show compact myocardium thickness in the left ventricle (lv) and right ventricle (rv). ivs, interventricular septum. **(B)** Transcriptome profiling of *Dll4^flox^;Cdh5-Cre^ERT^* hearts at E15.5. (a) left, Total number of differentially expressed genes identified by RNA-seq (BH adjusted P < 0.05; abs(logFC) > 0.5) in E15.5 *Dll4^flox^;Cdh5-Cre^ERT^*hearts. Numbers indicate upregulated genes (red) and downregulated genes (blue). Right, Circular plot showing representative differentially expressed genes associated to selected IPA functions belonging to the Cardiovascular System Development and Function category. Activation z-score scale: green repression; orange, activation; white, unchanged. LogFC scale: red, upregulated; blue, downregulated; white, unchanged. (b) Circular plot showing representative differentially expressed genes depending of selected upstream regulators. Colour codes for z-score and logFC scale as in (A). **(C)** Induced Dll4 deletion disrupts arterial differentiation. ISH for the arterial markers *Efnb2, Dll4*, and *Fabp4* on E15.5 heart sections from control and *Dll4^flox^;Cdh5-Cre*^ERT^mice tamoxifen-induced at E12.5. Scale bars, 100 µm.

**Supplemental Figure 7.**
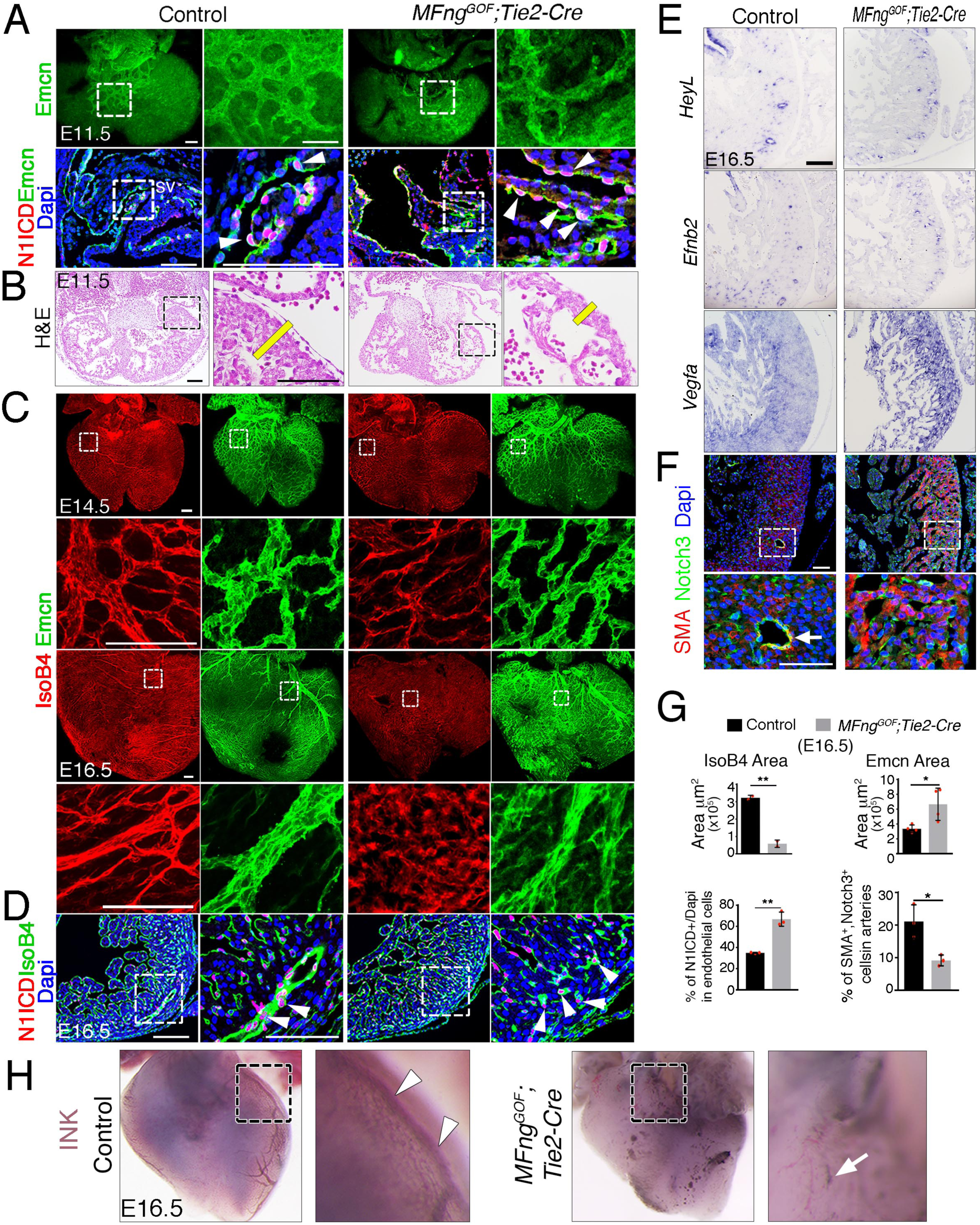
Forced *MFng* expression disrupts arterio-venous differentiation and arterial endothelial integrity. (**A**) *Top*, Dorsal view of whole-mount immunochemistry for Emcn (green) in E11.5 control and *MFng^GOF^;Tie2-Cre* hearts. *Bottom*, Immunohistochemistry for N1ICD (red) and Emcn (green) on E11.5 control and *MFng^GOF^;Tie2-Cre* heart sections. Dapi counterstain (blue). Arrowheads indicate N1ICD-stained nuclei. sv, sinus venosus. Scale bars, 100 µm. H&E staining of E11.5 control and *MFng^GOF^;Tie2-Cre* mutant heart sections. The yellow bars indicate compact myocardium (CM) thickness. Scale bars, 100 µm. (**B**) Dorsal view of whole-mount immunohstochemistry for IsoB4 (red) and Emcn (green) in E14.5 control and *MFng^GOF^;Tie2-Cre* hearts. (**C**) Dorsal view of whole-mount immunhistochemistry for IsoB4 (red) and Emcn (green) in E16.5 control and *MFng^GOF^;Tie2-Cre* hearts. Scale bars, 100 µm. (**D**) Immunohistochemistry for N1ICD (red) and IsoB4 (green) on E16.5 WT and *MFng^GOF^;Tie2-Cre* heart sections. Arrowheads indicate N1ICD-stained nuclei. Scale bars, 100 µm. (**E**) ISH of *Heyl, Efnb2*, and *Vegfa* on E16.5 control and *MFng^GOF^;Tie2-Cre* heart sections. Scale bar, 100 µm. (**F**) Immunohistochemistry for SMA (red) and Notch3 (green) on E16.5 control and *MFng^GOF^;Tie2-Cre* heart sections. Dapi counterstain (blue). Arrowhead indicates SMA-Notch3 co-immunostaining in coronary vessels. Scale bar, 50 µm. Microscope: Nikon A1-R. Software: NIS Elements AR 4.30.02. Build 1053 LO, 64 bits. Objectives: Plan Apo VC 20x/0,75 DIC N2 dry; Plan Fluor 40x/1,3 Oil DIC H N2 Oil. (**G**) Quantified data for E16.5 control and *MFng^GOF^;Tie2-Cre* hearts: area covered by coronary arteries; area covered by veins; percentage of N1ICD-positive nuclei in endothelium as a percentage (%) of total nuclei; and SMA-Notch3-co-immunostaining in coronary arteries. (**H**) Visualization of the coronary arterial vasculature at E16.5. Whole–mount aortic ink perfusion of the left coronary artery in control and *MFng^GOF^;Tie2-Cre* hearts. Note normal artery perfusion in the control (arrowheads) and the interrupted hemorrhaging artery in the transgenic heart (arrow). Data are mean ± s.d. (n= 3 sections from 3 control embryos and n = 3 from 3 transgenic embryos). \**P* < 0.05, \*\**P* < 0.01 by Student’s *t*-test.

**Supplemental Figure 8.**
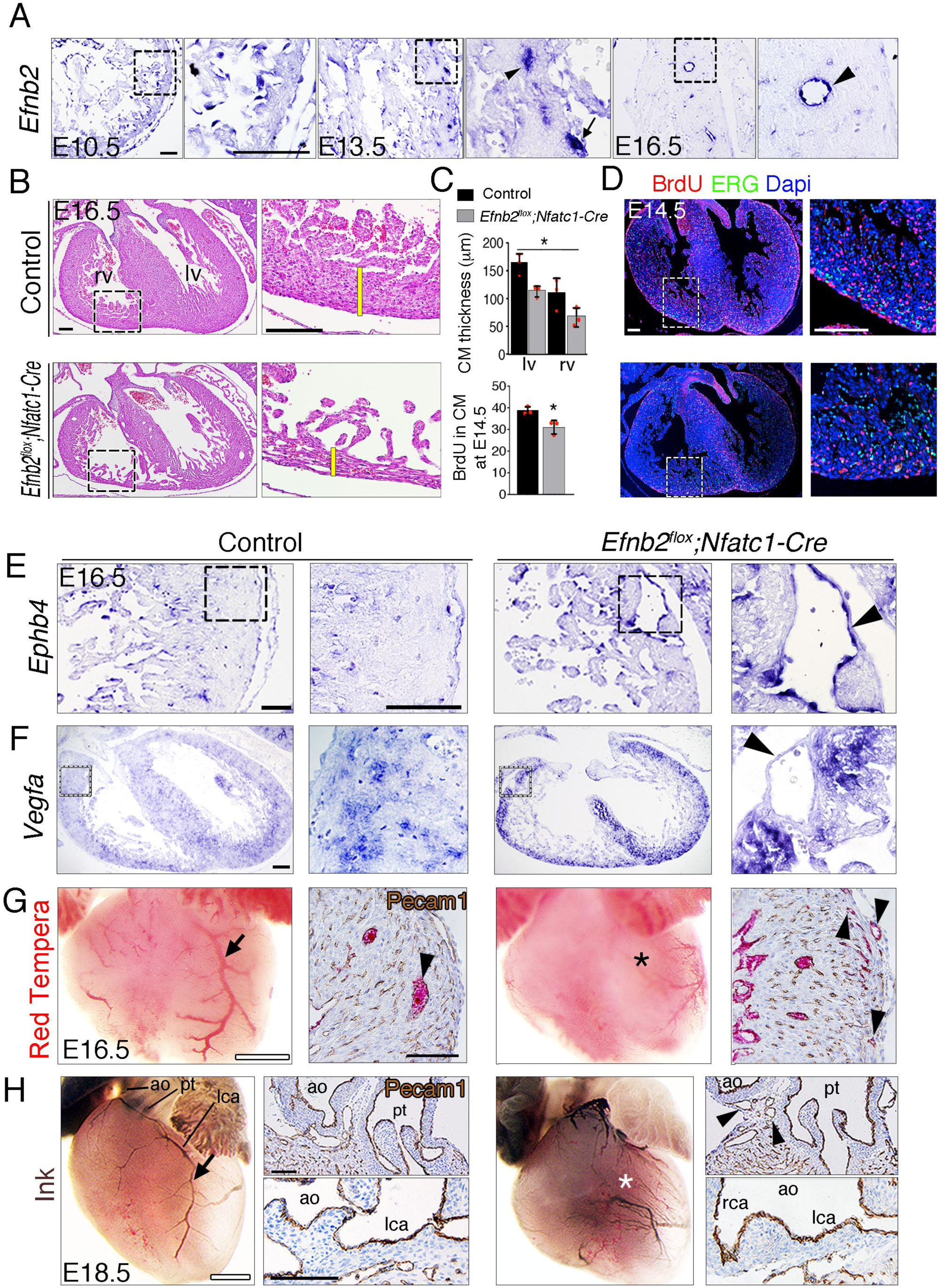
Endocardial *Efnb2* deletion disrupts myocardial proliferation, arterio-venous differentiation, and arterial endothelial integrity. **(A)** ISH in control embryos, showing *Efnb2* expression in chamber endocardium at E10.5 (arrowheads); coronary arteries (arrowheads) and a coronary vein (arrow) at E13.5; and a coronary artery at E16.5 (arrowhead). Scale bars, 50 µm. **(B)** H&E staining of transverse heart sections of E16.5 control and *Efnb2^flox^;Nfatc1-Cre* hearts. Yellow bars indicate compact myocardium (CM) thickness. BrdU (red) and ERG (green) staining in E14.5 control and *Efnb2^flox^;Nfatc1-Cre* hearts. Scale bars, 100 µm. (**C**) Charts showing (c) CM thickness in left ventricle (lv) and right ventricle (rv) and BrdU-stained nuclei in CM as a percentage (%) of all Dapi-stained nuclei. Data are mean +/-s.d. (n= 3 sections from 3 control embryos, and n=3 sections from 3 mutant embryos)**. (D)** BrdU (red) and ERG (green) staining in E14.5 control and *Efnb2^flox^;Nfatc1-Cre* hearts. Scale bars, 100 µm. Microscope: Nikon A1-R. Software: NIS Elements AR 4.30.02. Build 1053 LO, 64 bits. Objectives: Plan Apo VC 20x/0,75 DIC N2 dry; Plan Fluor 40x/1,3 Oil DIC H N2 Oil. **(E)** ISH of the venous marker *Ephb4* in E16.5 control and *Efnb2^flox^;Nfatc1-Cre* hearts, showing increased *Ephb4* expression and a severe arterio-venous malformation or fistula in the mutant (arrowhead). **(F)** ISH of *Vegfa* in E16.5 control and *Efnb2^flox^;Nfatc1-Cre* hearts, showing increased *Vegfa* expression in myocardium and an arterio-venous fistula (arrowhead). Scale bars, 50 µm. **(G)** Visualization of the coronary arterial vasculature at E16.5; whole-mount heart aortic Red Tempera perfusion reveals perfusion of the left coronary artery in the control (arrow) and the absence of perfusion in the *Efnb2^flox^;Nfatc1-Cre* mutant (asterisk). Scale bars, 500 µm. Pecam-1 staining of histological sections shows larger perfused coronaries in the control heart (arrowhead) and smaller perfused coronaries in the mutant (arrows). Scale bars, 100 µm. **(H)** Visualization of the coronary arterial vasculature at E18.5; whole-mount heart aortic ink perfusion reveals perfusion of the left coronary artery (lca) in control (arrow) and its absence in the *Efnb2^flox^;Nfatc1-Cre* mutant (asterisk). Scale bars, 500 µm. Pecam-1 staining of histological sections shows numerous peritruncal vessels surrounding the pulmonary trunk in the mutant (pt; arrowheads), while lca rooting to the aorta is maintained. Scale bars, 100 µm.

**Supplemental Figure 9.**
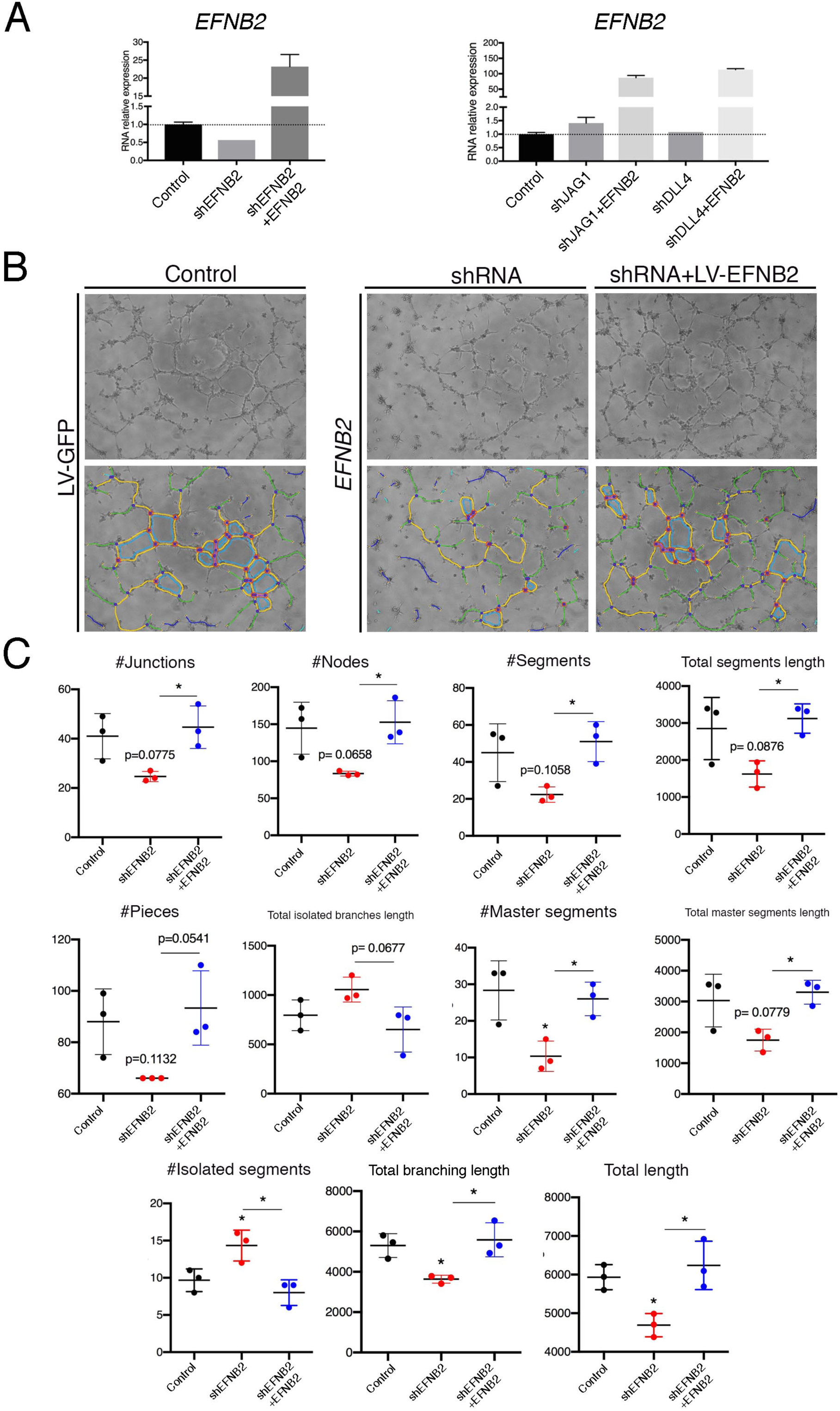
EPHRINB2 rescues defective capillary network formation after shRNA-mediated silencing of *EFNB2*. **(A)** qRT-PCR of expression of *EFNB2* after transduction of indicated shRNA. **(B)** Representative phase contrast images (1 of 2 experiments) of the HUVEC network, after transduction of indicated shRNA and rescue by *EFNB2* analyzed by the Angiogenesis Analyzer for ImageJ. **(C)** Informative measurements in the analyzed area: nodes surrounded by junctions (red surrounded by blue). Isolated elements (purple). Segments (yellow). Total segments length: sum of length of the segments. Number of pieces: sum of number of segments, isolated elements and branches detected. Total isolated branches length: sum of the length of the isolated elements. Red asterisks refer to comparisons between experimental and control situations. Black asterisks refer to comparisons between shRNA-mediated inhibition and LV-mediated rescue. Data are means ± s.d. \*\*\*\**P* < 0.0001; \*\*\**P* < 0.001; \*\**P* < 0.01; \**P* < 0.05, by ANOVA.

**Supplemental Table 1.** Excel file containing the lethality phases of *Jag1^flox^;Nfatc1-Cre, Dll4^flox^;Nfatc1-Cre, Jag1^flox^;Pdgfb-iCre^ERT2^, Dll4^flox^;Cdh5-iCre^ERT2^ Dll4 ^flox^;Cdh5-Cre^ERT2^, MFng^Gof^;Tie2-Cre, Efnb2 ^flox^;Nfatc1*-Cre embryos.

**Supplemental Table 2.** Excel file containing lists of differentially expressed genes identified after analysis of RNA-seq data, for contrasts between E12.5 *Jag1^flox^;Nfatc1-Cre* (Jag1Nfatc1 211Deg); E12.5 *Dll4^flox^;Nfatc1-Cre* (Dll4Nfatc1 274Deg); E16.5 *Dll4^flox^;Cdh5-Cre^ERT2^* (Dll4Cdh5 163Deg) and their control counterparts. Upregulated genes appear in red background and downregulated genes in blue.

**Supplemental Table 3.** Excel file describing results obtained with IPA for the sets of genes detected as differentially expressed (adjusted P value < 0.05) in contrasts between *Jag1^flox^;Nfatc1-Cre* (JAG1NF), *Dll4^flox^;Nfatc1-Cre* (Dll4NF), *Dll4^flox^;Cdh5-Cre^ERT2^* (Dll4Cdh5) and their control counterparts. Upstream Regulator analyses results, in sheets labelled as “UpRegTransc”, are restricted to molecules of type “transcription regulator”. For each predicted upstream regulator, tables describe expression log ratio (when available), predicted activation state, activation z-score, enrichment *P*_value and the collection of differentially expressed genes that are targets of the regulator. Downstream Effect analyses results, in sheets labelled as “CardSystDevFunc”, are restricted to functions of category “Cardiovascular System Development and Function”. For each functional term, tables describe enrichment *P* value, predicted activation state, activation z-score and the collection of differentially expressed genes associated to that particular function. Positive and negative z-score values suggest activation or inhibition of the corresponding upstream regulator or function in the mutant or control condition, respectively; abs(z-score) > 2 and *P* value < 0.05 are considered significant.

**Supplemental Table 4.** Excel file containing the list of primary and secondary antibodies used in this study to immunodetect proteins in whole-mount or paraffin sections.

